# PKCα-dependent interaction of otoferlin and calbindin: evidence for regulation of endocytosis in inner hair cells

**DOI:** 10.1101/779520

**Authors:** Andreia P. Cepeda, Hanan Al-Moyed, Christof Lenz, Henning Urlaub, Ellen Reisinger

**Affiliations:** Molecular Biology of Cochlear Neurotransmission Group, Department of Otorhinolaryngology, University Medical Center Göttingen, Göttingen, Germany; Collaborative Research Center 889, University of Göttingen, Göttingen, Germany; Göttingen Graduate Center for Neurosciences, Biophysics, and Molecular Biosciences, University of Göttingen, Göttingen, Germany; Core Facility Proteomics, Institute of Clinical Chemistry, University Medical Center Göttingen, Göttingen, Germany; Bioanalytical Mass Spectrometry Group, Max Planck Institute for Biophysical Chemistry, Göttingen, Germany

**Keywords:** calcium / inner ear / phosphorylation / ribbon synapse / synaptic transmission /

## Abstract

Otoferlin is essential for the fast and indefatigable release of synaptic vesicles at auditory inner hair cell (IHC) ribbon synapses, being involved in exocytic, endocytic and regenerative steps of the synaptic vesicle cycle. Serving diverse functions at this highly dynamic synapse implies that this multi-C_2_ domain protein is precisely regulated. Here we found protein kinase C α (PKCα) and otoferlin to colocalize in endocytic recycling compartments upon IHC depolarization and to interact in an activity-dependent manner. *In vitro* assays confirmed that PKCα can phosphorylate otoferlin at five serine residues, which correlates with increased serine phosphorylation in <40 nm proximity to otoferlin in murine IHCs that can be fully blocked by combining PKC and CaMKII inhibitors. Moreover, otoferlin interacts with calbindin-D28k in stimulated IHCs, which was precluded when PKCα was inhibited. Similarly, the activity-dependent increase in otoferlin-myosin VI interaction depends on PKCα activation. We propose that upon strong hair cell depolarization, PKCα phosphorylates otoferlin, thereby enabling it to interact with calbindin-D28k and myosin VI, building a Ca^2+^-dependent signaling complex that possibly regulates different modes of endocytosis.

## Introduction

In the mammalian auditory system, sound encoding between the sensory inner hair cells (IHCs) and the primary auditory neurons occurs with remarkable precision, reliability, and dynamics over prolonged periods of stimulation (Moser & Beutner, 2000; Glowatzki & Fuchs, 2002). IHC ribbon synapses are highly specialized for this challenging task, constantly sustaining the pool of fusion-competent vesicles. At physiological temperature, each synapse of a depolarized IHC can sustain a synaptic vesicle (SV) fusion rate of up to 2300 vesicles per second for at least several hundred milliseconds (Strenzke *et al*, 2016). This high release rate requires the efficient and coordinated retrieval of excess plasma membrane, for which both clathrin-independent and clathrin-dependent modes of endocytosis have been proposed (Neef *et al*, 2014; Kroll *et al*, 2019). Crucial proteins for Ca^2+^-triggered exocytosis in conventional synapses, like SNAREs, synaptotagmins, Munc13 or complexins, are either absent or are dispensable for exocytosis in IHCs (Safieddine & Wenthold, 1999; Strenzke *et al*, 2009; Beurg *et al*, 2010; Nouvian *et al*, 2011; Reisinger *et al*, 2011; Vogl *et al*, 2015). The multi-C_2_ domain protein otoferlin seems to replace some of these proteins and is currently hypothesized to act as the Ca^2+^ sensor for exocytosis in mature IHCs (Roux *et al*, 2006; Michalski *et al*, 2017). Different *OTOF* mutations lead to almost entirely abolished IHC exocytosis and thus to profound deafness in humans and animal models (Yasunaga *et al*, 1999, 2000; Roux *et al*, 2006; Longo-Guess *et al*, 2007; Marlin *et al*, 2010; Pangršič *et al*, 2010; Reisinger *et al*, 2011). Otoferlin is involved in vesicle priming and fusion, vesicle replenishment, vesicle reformation from bulk endosomes, active zone clearance, and clathrin-mediated endocytosis (Pangršič *et al*, 2010; Duncker *et al*, 2013; Jung *et al*, 2015; Strenzke *et al*, 2016). It has been reported to interact with several proteins involved in the SV cycle, e.g. Rab8b, myosin VI, Ca_V_1.3 calcium channels, the adaptor protein 2 (AP-2) and endophilin A (Roux *et al*, 2006; Heidrych *et al*, 2008, 2009; Ramakrishnan *et al*, 2009; Roux *et al*, 2009; Johnson & Chapman, 2010; Zak *et al*, 2012; Duncker *et al*, 2013; Ramakrishnan *et al*, 2014; Vincent *et al*, 2014; Jung *et al*, 2015; Hams *et al*, 2017; Meese *et al*, 2017; Kroll *et al*, 2019). Otoferlin bears six to seven C_2_ domains, of which at least three likely bind Ca^2+^ (Meese *et al*, 2017). Binding of Ca^2+^ to C_2_ domains is known to promote or hinder protein interactions, thus it is conceivable that this modular protein might integrate a series of regulatory interactions to finely balance the requirements of exo- and endocytosis at this synapse.

In conventional neuronal synapses, second messenger-activated protein kinases like Ca^2+^/calmodulin-dependent protein kinase II (CaMKII), cAMP-dependent protein kinase A (PKA), and protein kinase C (PKC) tightly and finely regulate synaptic transmission, and their activation correlates with increased transmitter release (Capogna *et al*, 1995; Hilfiker & Augustine, 1999). They control the function of the release machinery and the final steps of SV docking/fusion by regulating not only the availability of free SNARE proteins to form the functional fusion machinery but also protein-protein interactions within the release apparatus (reviewed in Turner *et al*, 1999; Leenders & Sheng, 2005). They have also been implicated in presynaptic plasticity via regulation of the refilling of the readily releasable pool of SVs thereby governing the number of release sites and the release probability (Pang *et al*, 2010; Stevens & Sullivan, 1998; Leenders & Sheng, 2005). At IHC synapses, CaMKIIδ was shown to phosphorylate otoferlin, rendering its C_2_F domain Ca^2+^-insensitive under physiological conditions (Meese *et al*, 2017). The regulation of otoferlin’s activity by CaMKIIδ may provide a molecular mechanism that influences the kinetics of exocytosis, endocytosis and vesicle replenishment in IHCs.

In this study, we assessed the effects of PKC in IHC synaptic function. Conventional PKCs (cPKCs; α, β and γ), the most abundant, structurally comprise a phospholipid-binding diacylglycerol (DAG)/phorbol ester-binding C_1_ domain, followed by a Ca^2+^-binding C_2_ domain and a C-terminal kinase moiety (reviewed in Rosse *et al*, 2010; Callender & Newton, 2017). They require Ca^2+^ and either membrane-bound DAG or phosphatidylserine (PS) for activation, but can also be activated by other DAG mimetics, resulting in enhanced glutamate release (Castagna *et al*, 1982; Nishizuka, 1984; Malenka *et al*, 1986; Shapira *et al*, 1987; Parfitt & Madison, 1993; Hori *et al*, 1999; Yawo, 1999; Brager *et al*, 2003; Korogod *et al*, 2007). The DAG/PKC pathway is one of the most potent pathways at presynaptic nerve terminals with its activation resulting in 50-100% potentiation of spontaneous and action potential-induced release (Malenka *et al*, 1986; Shapira *et al*, 1987; Lou *et al*, 2005). cPKC is autoinhibited by a pseudosubstrate sequence in its regulatory domain that sterically blocks the catalytic domain, rendering the kinase inactive and targeting it to the cytosol (Parker & Murray-Rust, 2004; Newton, 2010; Gould *et al*, 2011; Antal *et al*, 2014, 2015). cPKCs are activated in a sequential fashion. Firstly, Ca^2+^ binding to the C_2_ domain leads to an increased affinity of cPKC to phospholipids, resulting in its recruitment to membranes, where it then binds to its allosteric activator DAG via the C_1_ domain. This renders the cPKC in an open and active form ready to phosphorylate target substrates (Kraft *et al*, 1982; Sakai *et al*, 1997; Verdaguer *et al*, 1999; Sánchez-Bautista *et al*, 2006; Evans *et al*, 2006). Besides phosphorylating presynaptic proteins to facilitate exocytosis (Shimazaki *et al*, 1996; Kataoka *et al*, 2000; Genoud *et al*, 2001; Nagy *et al*, 2002; Barclay *et al*, 2003; Wierda *et al*, 2007; Genç *et al*, 2014; Cijsouw *et al*, 2014; Jong *et al*, 2016), PKC is emerging as a central player in vesicular transport pathways, being involved in regulation of membrane trafficking and endocytosis (reviewed in Alvi *et al*, 2007). For instance, it was shown to regulate the targeting of synaptotagmin IX to endocytic compartments (Haberman *et al*, 2005).

In this study, we investigated the effects of PKCα activation on the function of the IHC synaptic protein otoferlin, unravelling a Ca^2+^-controlled signalling complex potentially acting in regulation of endocytosis.

## Results

### PKCα is expressed in the organ of Corti and redistributes to the synaptic region of IHCs upon stimulation where it colocalizes with otoferlin

PKC is known to regulate presynaptic plasticity, exocytosis and endocytosis in neurons (Shapira *et al*, 1987; Alvi *et al*, 2007; Jong *et al*, 2016). To examine whether PKC is involved in modulating presynaptic function in IHCs, we mapped its subcellular localization and tested if it colocalizes with otoferlin (Figs 1 and EV1). We performed immunohistochemistry on acutely dissected organs of Corti from wild-type C57BL/6J (henceforth, WT) mice after the onset of hearing (at P15). Dissections were done in phosphate buffered saline (PBS) solution (no supplemented Ca^2+^). The α isoform was chosen because it is the cPKC most ubiquitously expressed in other systems and there is only one variant (Kofler et al, 2002). We found PKCα to be expressed both in IHCs and OHCs (Fig EV1A). In IHCs, PKCα was expressed throughout the cell (Fig EV1B), predominantly in the cytosol and to a lesser extent at the plasma membrane, where it partially colocalizes with otoferlin (Fig EV1C-D).

Upon Ca^2+^ influx, PKC typically relocates to the plasma membrane (Codazzi *et al*, 2001; Zhao *et al*, 2006). Using a previously described rest/stimulation/recovery paradigm (Kamin *et al*, 2014; Revelo *et al*, 2014), we followed PKCα immunofluorescence in IHCs (Fig 1). In resting conditions (1 min, 5.36 mM KCl, no supplemented Ca^2+^), PKCα immunoreactivity was located almost exclusively in the cytosol. Strong stimulation (1 min, 65.36 mM KCl, 2 mM CaCl_2_) resulted in a distinct relocation of PKCαwithin the base of the IHCs where it accumulated in clusters close to the plasma membrane (Figs 1A-B and 2). Most PKCα clusters were found near the synaptic ribbons (Fig 2A-F) in structures resembling plasma membrane patches and endosomes where it partially colocalized with otoferlin (Fig 2E-H). This effect seems to be rather transitory as only few PKCα clusters remained after 5-minute stimulation, and hardly any PKCαclusters persisted after 5-minute recovery (5 min, 5.36 mM KCl, 2 mM CaCl_2_, following a 1-minute stimulation, Fig 1A-B). The relocation of PKCα to regions close to active zones in a cluster-like appearance seems to occur only for strong stimulations, as no clustering could be observed at milder stimulations inducing less Ca^2+^ influx (1 min, 25 mM KCl, 2 mM CaCl_2_; Fig EV2A). Interestingly, the trafficking of otoferlin seems to follow that of PKCα (Fig 1A, C-E), pointing towards a probable interaction of the two proteins. At rest, otoferlin immunofluorescence was found throughout the cell in the cytoplasm and at the plasma membrane whereas PKCα seems to be expressed mainly in the cytoplasm with rather weak plasma membrane localization (Fig 1C, upper panel and 1D, apical/basal PKCα ratio: 1.05±0.03; mean ± standard error of the mean, s.e.m.). After 1-minute stimulation, both PKCα and otoferlin localized more towards the basal region of the IHCs when compared to resting conditions (Fig 1C, middle panel and Fig 1D, apical/basal ratio <1), while both were found more apically after letting the cells recover for 5 minutes (Fig 1C, bottom panel and Fig 1D, apical/basal ratio >1).

**Figure 1.**
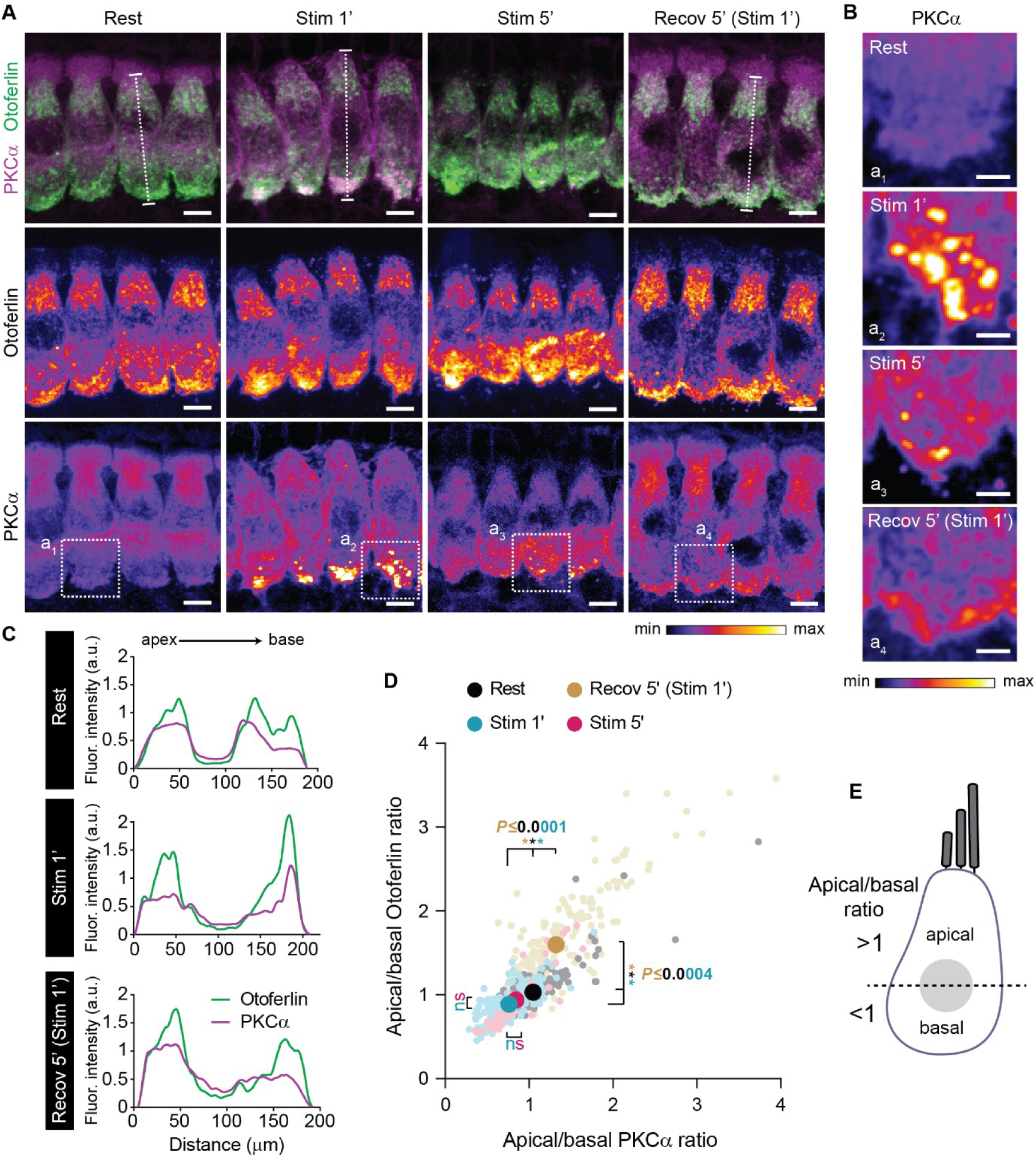
PKCα redistributes to base of IHCs upon strong stimulation and PKCα distribution correlates with otoferlin localization. **A** High magnification views of representative WT P14-16 IHCs immunolabeled for PKCα and otoferlin, at rest (Rest), after strong stimulation for 1 (Stim 1’) and 5 minutes (Stim 5’) and after a 5-minute recovery period post 1-minute stimulation (Recov 5’ (Stim 1’)). For clarity, individual otoferlin and PKCα channels are depicted separately with an intensity-coded lookup table with warmer colors representing higher pixel intensities (middle and bottom panels). **B** PKCα immunolabelling (intensity-coded lookup table) in higher magnification views of basal regions of the IHCs labelled in (A) as a1, a2, a3, a4. Note that PKCαclustering is maximal at 1-minute stimulation. **C** Fluorescence intensity line profile through the longitudinal axis at the mid-region of representative IHCs labelled in (A), from apex to base (five optical sections). Note that PKCα relocates to the base of the IHCs at Stim 1’ where the intensity line profile overlaps with that of otoferlin. **D** Correlation of otoferlin and PKCα ratio of apical/basal immunofluorescence (above/below nuclear midline depicted in (E)) indicates a strong localization correlation between PKCα and otoferlin. Mean values are displayed with darker colors and bigger symbols. Individual cells are depicted with lighter colors and smaller symbols. Mean averages, sample size and statistical analysis are detailed in *Appendix Table S1*. **E** Schematic representation of an IHC, indicating how the apical/basal ratio of immunofluorescence was determined. Higher immunofluorescence above or below the nuclear midline (dashed line) results in a ratio >1 or <1, respectively. A ratio >1 indicates a shift towards a more apical localization while a ratio <1 corresponds to a more basal localization of the protein. Data information: In (A-B), maximum intensity projections of confocal optical sections. Scale bars: 5 μm (A), 2 μm (B). Fluor., fluorescence.

**Figure 2.**
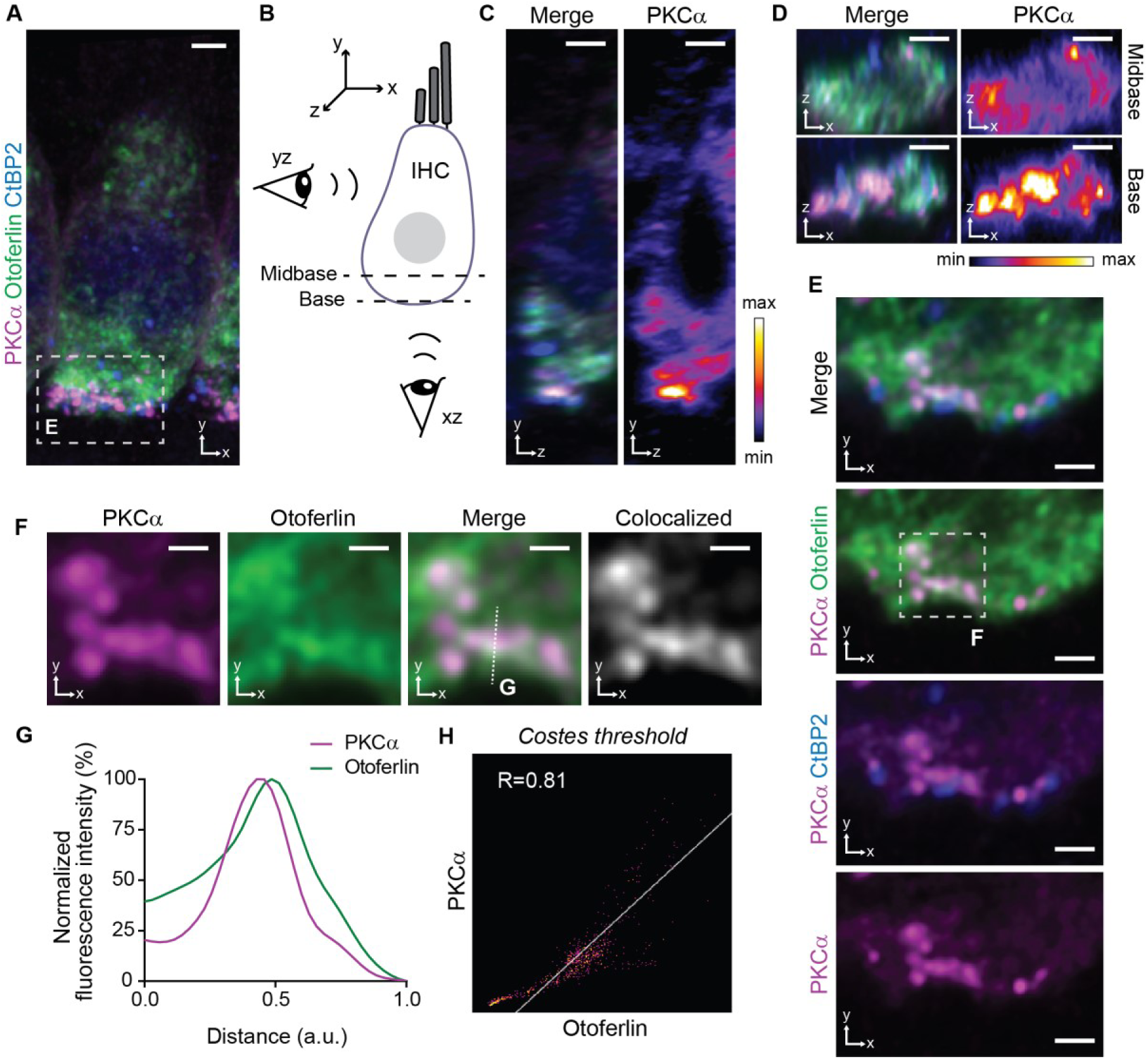
PKCα redistributes to the base of IHCs after strong stimulation, where it is found near the synaptic ribbons and partially colocalizes with otoferlin. **A** High magnification view of a representative WT P15 IHC immunolabeled for PKCα (magenta), otoferlin (green) and the ribbon marker CtBP2 (blue), after strong stimulation for 1 minute. **B-D** Orthogonal views of the IHC displayed in (A). (B) Schematic representation of an IHC, with illustration of side (yz; C) and bottom (xz; D) views, for clarity. In (C-D), individual PKCα channel is depicted separately (right panels) with an intensity-coded lookup table. **E** Higher magnification view of the basal region of the IHC displayed in (A), showing PKCα accumulations in regions close to the ribbons (immunolabelled with an antibody against CtBP2, blue) and in endosome-like structures. **F-H** Colocalization analysis for otoferlin (green) and PKCα (magenta) channels for the area labelled in E. (F) Single image plane with individual (first two panels) and merged (third panel) channels. Pixels with positive signals for both channels are shown in white (forth panel). (G) PKCα and otoferlin intensity profiles through the dashed line in F. (H) Scatterplot of PKCα and otoferlin pixel intensities and calculated Pierson’s correlation coefficient using the Costes automatic thresholding method. Data information: In (A), maximum intensity projection of confocal optical sections. In (C-F), single confocal optical sections. Scale bars: 2 μm (A, C-D), 1 μm (E), 0.5 μm (F). IHC, inner hair cell.

In order to find out if the observed clustering of PKCα close to the plasma membrane of IHCs is coherent with its activation, we assessed its localization after pharmacological PKCα activation. In many cell types treatment with phorbol 12-myristate 13-acetate (PMA), a PKC agonist that mimics DAG and strongly binds cPKCs (Takekoshi *et al*, 1995), induced the recruitment of PKC to membranes (Hermelin *et al*, 1988; Huang *et al*, 1997; Feng *et al*, 1998, 2000; Tardif *et al*, 2002; González *et al*, 2003; Schechtman *et al*, 2004; Wu *et al*, 2006; Cordey & Pike, 2006). When we treated organs of Corti with PMA, PKCα redistributed to the plasma membrane throughout the cell (Fig EV2B-C) with the stronger effect observed at 15-minute incubation (Fig EV2B-C, PMA 15’). Moreover, PMA treatment for 5 minutes resulted in enrichment of otoferlin immunofluorescence in large patches at the base of the hair cells (Fig EV2B, PMA 5’) and enhanced colocalization of PKCα and otoferlin at the basolateral plasma membrane could be observed for 15-minute incubation (Fig EV2B, PMA 15’).

Altogether, these results indicate co-trafficking of PKCα and otoferlin upon PKCα activation (either pharmacologically or following strong stimulation) and thus point towards a possible activity-dependent interaction of the two proteins.

### PKCα interacts with otoferlin in IHCs

We next investigated a potential interaction of PKCα and otoferlin in IHCs. We first performed a proximity ligation assay (PLA), which allows *in situ* detection of endogenous protein interactions with single molecule resolution, detecting a <40 nm distance of antibody-labeled proteins (Fig 3A,B). This assay was previously established for rat IHCs with the reported interaction pair otoferlin-myosin VI and was validated here in mouse IHCs (Appendix Fig S1). The PLA for otoferlin and PKCα performed in explanted organs of Corti of P14-16 mice in resting conditions resulted in few fluorescent puncta distributed throughout the IHC (Fig 3 A,B). When the same PLA was performed after 1-minute stimulation, we saw a >4-fold increase in PLA fluorescence intensity (442±28%, n=141 IHCs) when compared to resting conditions (100±7%, n=122 IHCs), pointing to an interaction of the proteins upon strong IHC stimulation. The intensity of the PLA puncta dropped to 178±7% (n=112 IHCs) during a 5-minute recovery period, indicating a rather short-living otoferlin-PKCα complex (Fig 3A, B).

**Figure 3.**
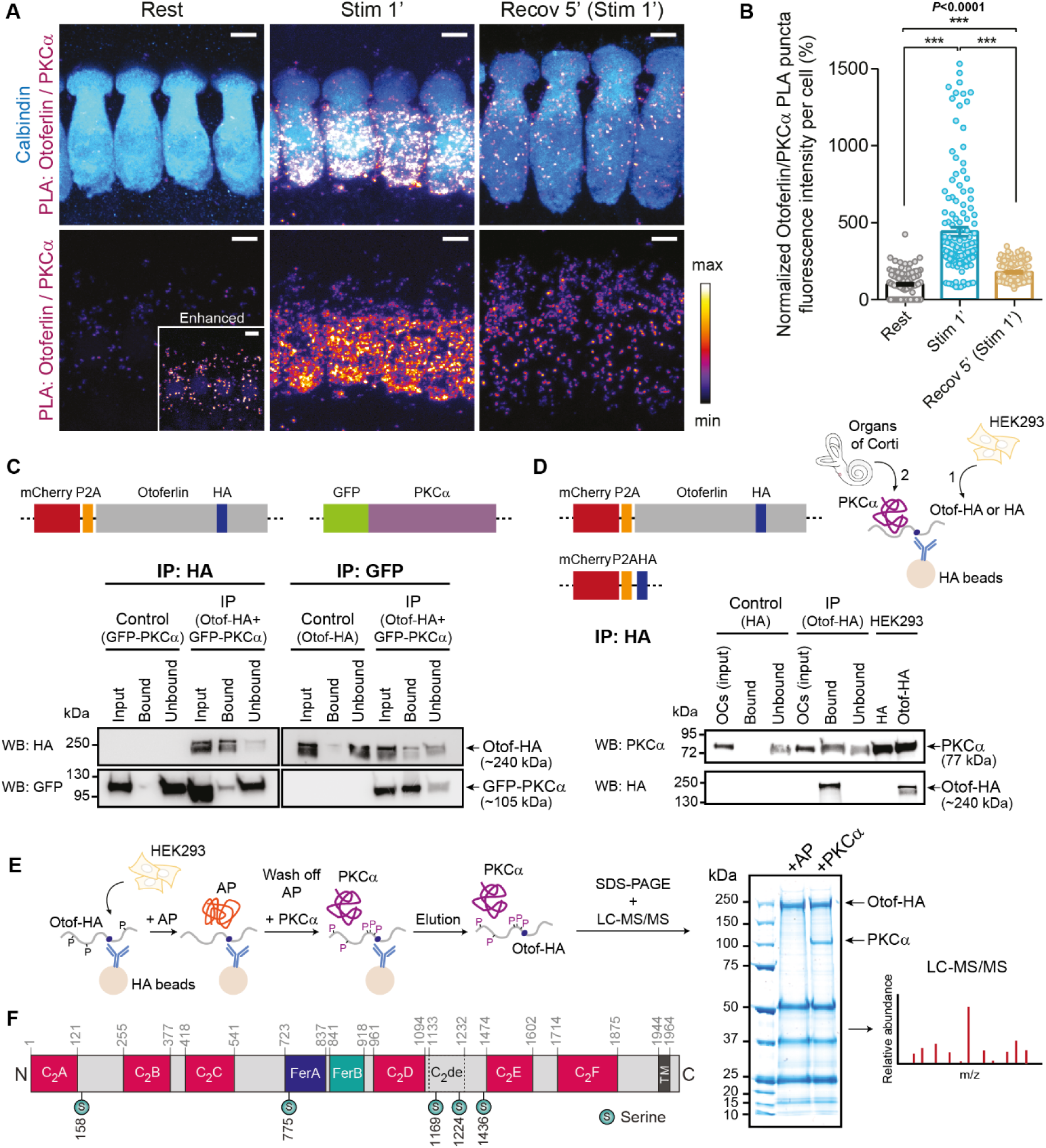
PKCα and otoferlin interact in IHCs. **A, B** PLA for otoferlin and PKCα performed on WT P15 IHCs at rest, after strong stimulation for 1 (Stim 1’) and 5 minutes (Stim 5’), and after a 5-minute recovery period post stimulation (Recov 5’ (Stim 1’)). (A) High magnification views of representative PLAs: calbindin (blue) was used as IHC marker; PLA channel is depicted with an intensity-coded lookup table with warmer colors representing higher pixel intensities. (B) Average otoferlin/PKCα PLA puncta fluorescence intensity per cell for all conditions, normalized to the resting condition. Individual cells are depicted with lighter colors and open symbols. See *Appendix Figure S2A and C* for control PLAs. **C** Representative immunoblot showing results of anti-HA and anti-GFP co-immunoprecipitation from lysates of HEK293 cells co-transfected with otoferlin-HA and GFP-PKCα. Samples were probed for HA and GFP. Upper panel depicts constructs used in binding assays. **D** Representative immunoblot showing results from pull-down assay from organs of Corti loaded onto anti-HA beads with previously bound otoferlin-HA expressed in HEK293 cells. Samples were probed for HA and PKCα. Upper left panel depicts constructs used in binding assays. Upper right panel depicts scheme of the assay. **E** Schematic representation of the *in vitro* phosphorylation assay and subsequent mass spectrometry analysis to assess PKCα-induced phosphosites on otoferlin. Alkaline phosphatase (AP) was used to remove any residual phosphate groups and obtain dephosphorylated otoferlin. After incubation with AP + PKCα or with AP only, samples were loaded onto a SDS-PAGE gel, otoferlin bands were excised, digested and analyzed by LC-MS/MS. For annotated MS/MS spectra of detected phosphosites, LC-MS/MS profiling of phosphopeptides and mapping of sites in the otoferlin sequence refer to *Appendix Figures S3-S9*. **F** Position of the phosphorylation sites in the otoferlin sequence (mouse, isoform 4, NP_001300696.1) determined by LC-MS/MS. For mapping of sites in the otoferlin sequence refer to *Appendix Figure S10*. Data information: In (A), maximum intensity projections of confocal optical sections. Scale bars: 5 μm. In (B), data are displayed as mean ± s.e.m.; ***P≤0.001 (Kruskal-Wallis test followed by Dunn’s multiple comparison test); mean averages, sample size and statistical analysis are detailed in *Appendix Table S1*. In (A-B), only puncta inside the cells were considered for quantification purposes; quantification was performed in *Imaris* as described in *Materials and Methods*. PLA, proximity ligation assay. IHC, inner hair cell. Otof, otoferlin.

Given that a positive PLA signal could potentially result from an indirect interaction via scaffolding proteins, we assessed whether otoferlin and PKCα interact directly *in vitro* (Fig 3C, D). In a first approach, we co-transfected HEK293T cells with HA-tagged full-length otoferlin (mCherry-P2A-mOtof-HA) and GFP-tagged PKCα (eGFP-PKCα; Fig 3C, upper panel) and performed anti-HA and anti-GFP co-immunoprecipitation (Co-IP) assays. Western blotting of immunoprecipitated samples of HA IPs, where otoferlin-HA was used as bait, revealed a band of ∼105 kDa in the eluate corresponding to GFP-PKCα when immunoblotted against GFP (Fig 3C, bottom left panel). Conversely, a band of ∼240 kDa corresponding to otoferlin-HA was detected with an anti-HA antibody, when GFP-PKCα was used as bait in GFP IPs. The faint bands in both situations suggest a weak interaction of the two proteins, possibly because the cells were harvested in conditions with weak PKC activation. In a second approach, we ran pull-down assays from organ of Corti homogenates which we loaded onto HA beads enriched with otoferlin-HA protein previously expressed in HEK293T cells (Fig 3D, upper panel). When we immunoblotted for PKCα, a strong band of ∼77 kDa was evident in the eluate; the same band was absent in control experiments with HEK293T-expressed HA peptide (Fig 3D, bottom panel). Both *in vitro* assays indicate that otoferlin and PKCα can interact directly.

An *in vitro* assay, where immobilized otoferlin-HA was incubated with recombinant PKCα was conducted to test if PKCα can phosphorylate otoferlin (Fig 3E). LC-MS/MS analysis (Appendix Figs S3 to S10) revealed phosphorylation of otoferlin at five serine residues: S158, S775, S1169, S1224 and S1436 (otoferlin variant 4, NP_001300696.1, Fig 3F). All phosphorylation sites were found to be conserved between mammalian and non-mammalian otoferlin orthologs, with the exception of S1169 at the C_2_de domain conserved only among mammalian species (Appendix Fig S11). Interestingly, none of the phosphorylated serine residues is located in one of the main six C_2_ domains; yet, two were found in the C_2_de domain, a putative C_2_ domain with poor conservation of sequence and secondary structure elements among species. Phosphorylation at S775 in the FerA domain could possibly alter the interaction of FerA with membranes in the presence of Ca^2+^ (Harsini *et al*, 2018). It is noteworthy that three of the five positions (S158, S775, S1224) match phosphorylation sites retrieved by different kinase-specific, sequence- and structure-based prediction tools (Appendix Fig S12 and Table S2).

In the absence of otoferlin, hardly any membrane turnover takes place in IHCs due to the abolishment of fast exocytosis (Roux *et al*, 2006). Here, we used knock-out mice (*Otof^−/−^*) (Reisinger *et al*, 2011) to find out if the disaggregation of PKCα immunofluorescence clusters depends on proper endocytic and vesicle recycling processes (Fig 4). In *Otof^−/−^* IHCs, although PKCα immunofluorescence levels were only slightly altered when compared to WT IHCs (WT: 100±2%, n=233 IHCs vs. *Otof^−/−^*: 92±1%, n=205 IHCs; *\*\*P=*0.0054, Mann-Whitney two-tailed t-test), the localization of PKCα was shifted towards the base of the cell (apical/basal ratio: WT 1.06±0.03 vs. *Otof^−/−^* 0.66±0.02; Fig 4A-C). Upon high potassium (K^+^) stimulation, PKCα relocated to the base of *Otof^−/−^* IHCs (Fig 4D-G), in a similar fashion to what happened in WT IHCs (Fig 1A-D). This indicates that exocytosis seems not to be required for the trafficking of PKCα towards the plasma membrane. However, after 5-minute stimulation and during recovery (following 1-minute stimulation), PKCα immunofluorescence was still evident at endosomal structures and at the basolateral plasma membrane in *Otof^−/−^* IHCs (Fig 4D-E). Given that otoferlin is not only involved in exo-endocytosis coupling but is also required for proper vesicle reformation from recycling endosomes, it is probable that these plasma membrane patches and endosomes are not converted to smaller vesicles as quickly as in the presence of otoferlin, reassuring that PKCα indeed persists in endosomal compartments in *Otof^−/−^* IHCs.

**Figure 4.**
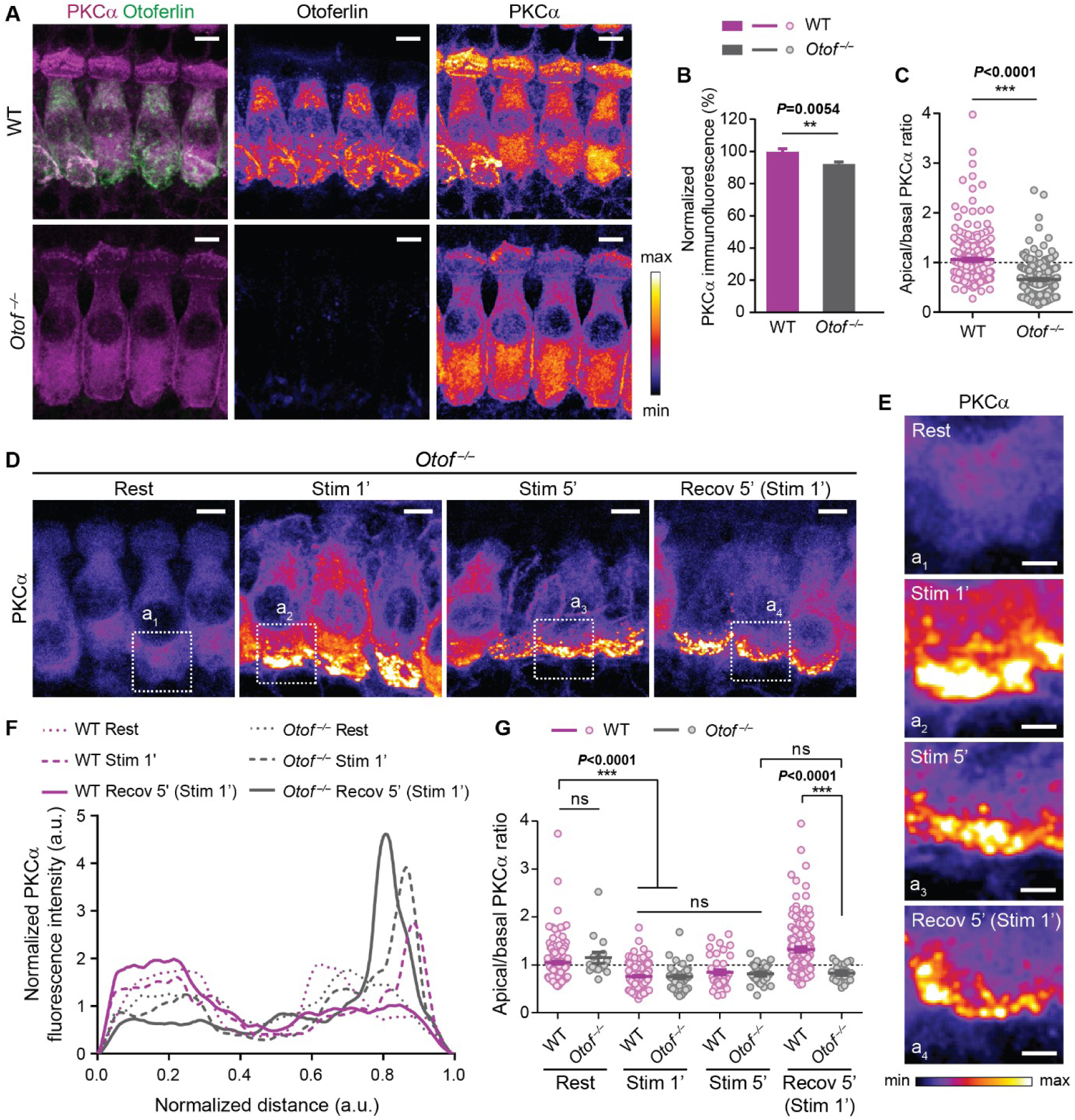
PKCα subcellular distribution is affected in otoferlin knock-out IHCs. **A-C** PKCα immunofluorescence in *Otof^−/−^* compared to WT P15-16 IHCs. (A) High magnification views of representative WT and *Otof^−/−^* P14-16 IHCs immunolabeled for PKCα (magenta) and otoferlin (magenta). Otoferlin and PKCα channels are depicted separately with an intensity-coded lookup table with warmer colors representing higher pixel intensities. (B) Quantification of overall PKCα immunofluorescence. (C) Apical/basal PKCα immunofluorescence (above/below nuclear midline). Numbers of cells in (C) also apply to (B). **D, E** PKCα distribution in *Otof^−/−^* P14-16 IHCs for all indicated conditions. (D) High magnification views of representative *Otof^−/−^* P15-16 IHCs immunolabeled for PKCα. (B) Higher magnification views of basal regions of the *Otof^−/−^* IHCs labelled in (D) as a1, a2, a3, a4. For clarity, an intensity-coded lookup table was used. **F, G** Comparison of PKCα distribution in WT B6 and *Otof^−/−^* P14-16 IHCs. (F) Fluorescence intensity line profile through the longitudinal axis at the mid-region of representative WT B6 and *Otof^−/−^* IHCs, from apex to base (five optical sections). (G) Comparison of apical/basal PKCα immunofluorescence (above/below nuclear midline) among the different experimental conditions. Data information: In (A, D-E), maximum intensity projections of confocal optical sections. Scale bars: 5 μm (A, D), 2 μm (E). In (B-C, G), data are displayed as mean ± s.e.m.; ns P>0.05, **P≤0.01, ***P≤0.001; mean averages, sample size and statistical analysis are detailed in *Appendix Table S1*. In (C, G) individual cells are depicted with lighter colors and open symbols. Rest, resting; Stim 1’, 1-minute stimulation; Stim 5’, 5-minute stimulation; Recov 5’ (Stim 1’), 5-minute recovery after 1-minute stimulation. IHC, inner hair cell.

### Activity-dependent phosphorylation of otoferlin or otoferlin interactors by PKCα

To test whether otoferlin and/or proteins interacting with otoferlin are phosphorylated and to assess if the phosphorylation of otoferlin is activity-dependent *in vivo*, Meese *et al* (2017) applied a PLA to find phosphoserine residues in <40 nm distance from otoferlin in rat IHCs. Upon stimulation with high K^+^ the PLA signal increased when compared to resting conditions, and this effect could be only partially blocked by the CaMKII inhibitor KN-93 (Meese *et al*, 2017; Fig 10B-C), suggesting the involvement of other kinases in the regulation of synaptic function through phosphorylation of otoferlin in mammalian IHCs.

We sought to assess whether the increase in phosphorylation of otoferlin or otoferlin-associated proteins is complemented by PKC in mouse IHCs (Fig 5). We first stimulated the cells (with 65.36 mM KCl, 2 mM CaCl_2_) and observed an increase in PLA signal that peaked at 5-minute stimulation (Rest: 100±11%, n=100 IHCs vs. Stimulation 1’: 234±13%, n=37 IHCs vs. Stimulation 5’: 438±38%, n=52 IHCs; \*\*\**P*<0.0001, Kruskal-Wallis test followed by Dunn’s multiple comparison test; Fig 5). Pre-incubation with the PKC inhibitor bisindolylmaleimide I (BIM I) for 15 minutes prior to 5-minute stimulation blocked the stimulation-dependent increase in PLA signal to a large extent (BIM I + Stimulation 5’: 122±2%, n=34 IHCs; \*\**P*=0.0013 vs. Stimulation 5’). Moreover, pre-incubation with BIM I and KN-93 completely blocked this effect (BIM I+KN-93 + Stimulation 5’: 97±4%, n=50 IHCs; \*\*\**P*<0.0001; Fig 5). Treatment with the PKCα activator PMA led to an increase in PLA signal (PMA 5’: 157±4%, n=66 IHCs and PMA 15’: 139±6%, n=48 IHCs; \*\*\**P*≤0.0002 vs. Rest). This effect could only be seen for longer incubations times (PMA 1’: 77±7%, n=61 IHCs; ns *P*=0.3464 vs. Rest; Fig 5) likely due to extracellular application and thus the time needed for PMA to flip to the inner leaflet of the lipid membrane. The PMA-induced increase in PLA signal was not as evident as for 1- and 5-minute high K^+^ stimulations. Since PMA strongly activates PKCα even in the absence of Ca^2+^, the finding that phosphorylation by PKCα is enhanced under depolarizing conditions suggests that Ca^2+^ might be able to strengthen the interaction of otoferlin and PKCα, likely via binding to their C_2_ domains. Altogether, these data indicate that the activity-dependent phosphorylation of otoferlin and/or its interactors in mouse IHCs relies on the combined action of PKCα and CaMKIIδ.

**Figure 5.**
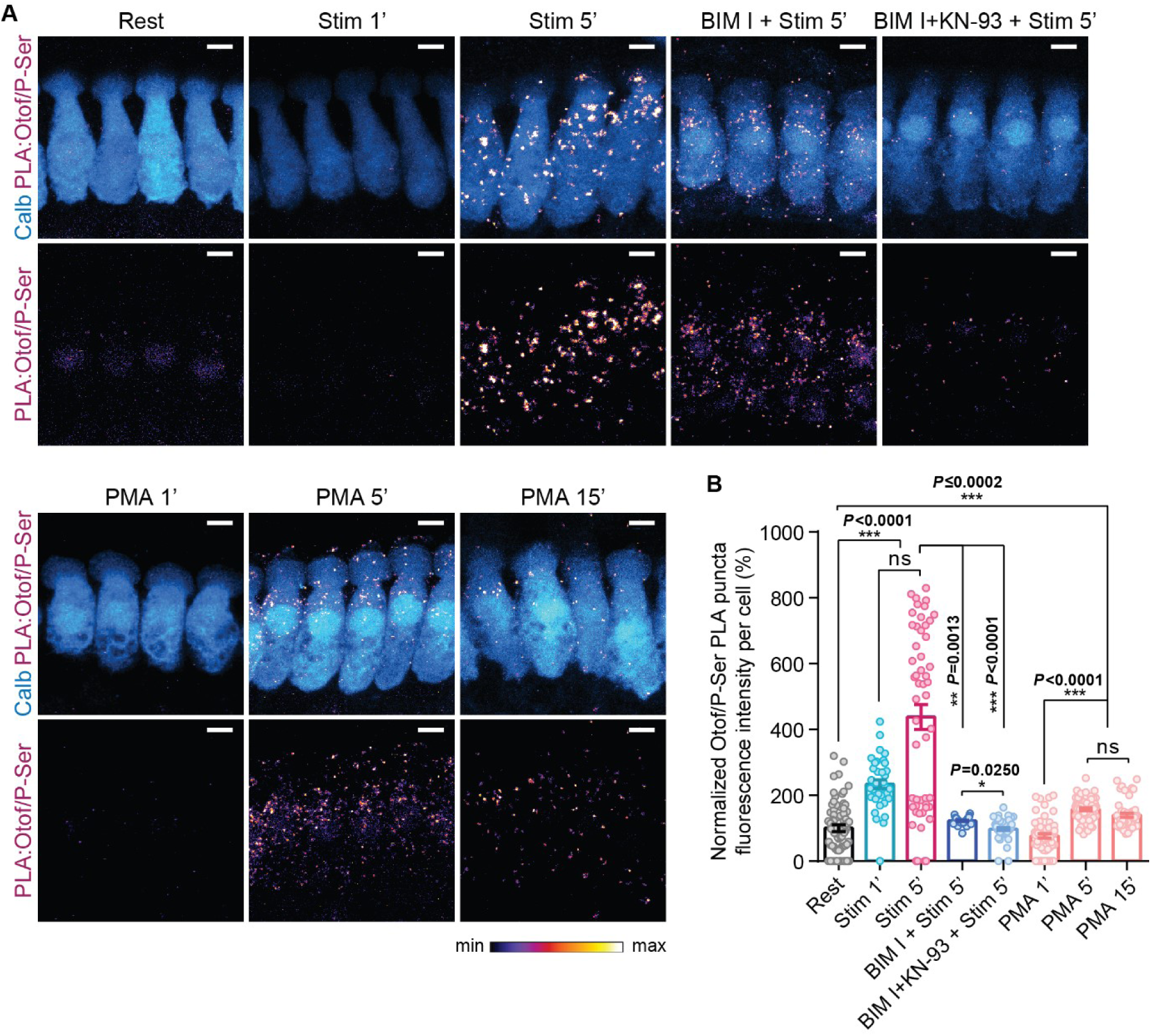
Otoferlin phosphorylation is strongly promoted by hair cell stimulation and can be blocked by combined inhibition of PKC and CaMKII. **A, B** PLA for otoferlin and phosphoserine residues performed on WT P15-16 IHCs at rest, after stimulation, after incubation with BIM I (PKC inhibitor) or with BIM I and KN-93 (CaMKII inhibitor), and after incubation with PMA (PKC activator). (A) High magnification views of representative PLAs; Calbindin (blue) was used as IHC marker; PLA channel is depicted with an intensity-coded lookup table with warmer colors representing higher pixel intensities. (B) Average otoferlin/phosphoserine PLA puncta fluorescence intensity per cell, normalized to the resting condition. Individual cells are depicted with lighter colors and open symbols. See control PLAs in *Appendix Figure S2A and B*. Data information: In (A), maximum intensity projections of confocal optical sections. Scale bars: 5 μm. In (B), data are displayed as mean ± s.e.m.; ns *P*>0.05, \**P*≤0.05, \*\**P*≤0.01, \*\*\**P*≤0.001 (Kruskal-Wallis test followed by Dunn’s multiple comparison test); mean averages, sample size and statistical analysis are detailed in *Appendix Table S1*. Rest, resting; Stim 1’, 1-minute stimulation; Stim 5’, 5-minute stimulation; BIM I + Stim 5’, Incubation with BIM I prior to 5-minute stimulation; BIM I+KN-93 + Stim 5’, Incubation with BIM I and KN-93 prior to 5-minute stimulation; PMA 1’, 1-minute incubation with PMA; PMA 5’, 5-minute incubation with PMA; PMA 15’, 15-minute incubation with PMA. IHC, inner hair cell. Otof, otoferlin. P-Ser, phosphoserine. Calb, calbindin. PLA, proximity ligation assay.

### Otoferlin interacts with myosin VI in a PKCα-dependent manner, but not with Vglut3

To narrow down the cellular pathways potentially affected by the PKCα-mediated regulation of otoferlin, we tested an activity-dependent interaction of otoferlin with candidate proteins.

Firstly, we assessed if the interaction of otoferlin with its reported interaction partner myosin VI (Heidrych *et al*, 2009; Roux *et al*, 2009) is influenced by IHC stimulation and PKC activation (Fig 6 A-C). After stimulation with high K^+^ for 1 minute, not only did myosin VI immunofluorescence levels increase significantly (Rest: 100± 5%, n=43 IHCs vs. Stimulation 1’: 138±3%, n=44 IHCs; \*\*\**P*<0.0001; Fig 6A-B) but the PLA signal for otoferlin and myosin VI increased even more (Rest: 100±3%, n=265 IHCs vs. Stimulation 1’: 173±4%, n=170 IHCs; *P*<0.0001; Fig 6C-D). The PLA signal remained high during a 5-minute recovery period following the 1-minute stimulation, indicating that the interaction persists in this time frame (Recovery 5’ Stimulation 1’: 183±4%, n=153 IHCs; *P*>0.9999 vs. Stimulation 1’). Incubation with BIM I prior to 1-minute high K^+^ stimulation led to a complete abolishment of the stimulation-induced increase in PLA signal (BIM I + Stimulation 1’: 102±6%, n=37 IHCs). Treatment with PMA also resulted in an increase in PLA signal (PMA 5’: 133±3%, n=122 IHCs and PMA 15’: 149±9%, n=96 IHCs; \*\*\**P*<0.0001 vs. Rest; Kruskal-Wallis test followed by Dunn’s multiple comparison test; Fig 6C-D). These results support the notion that the interaction of otoferlin and myosin VI is strongly PKC-dependent.

**Figure 6.**
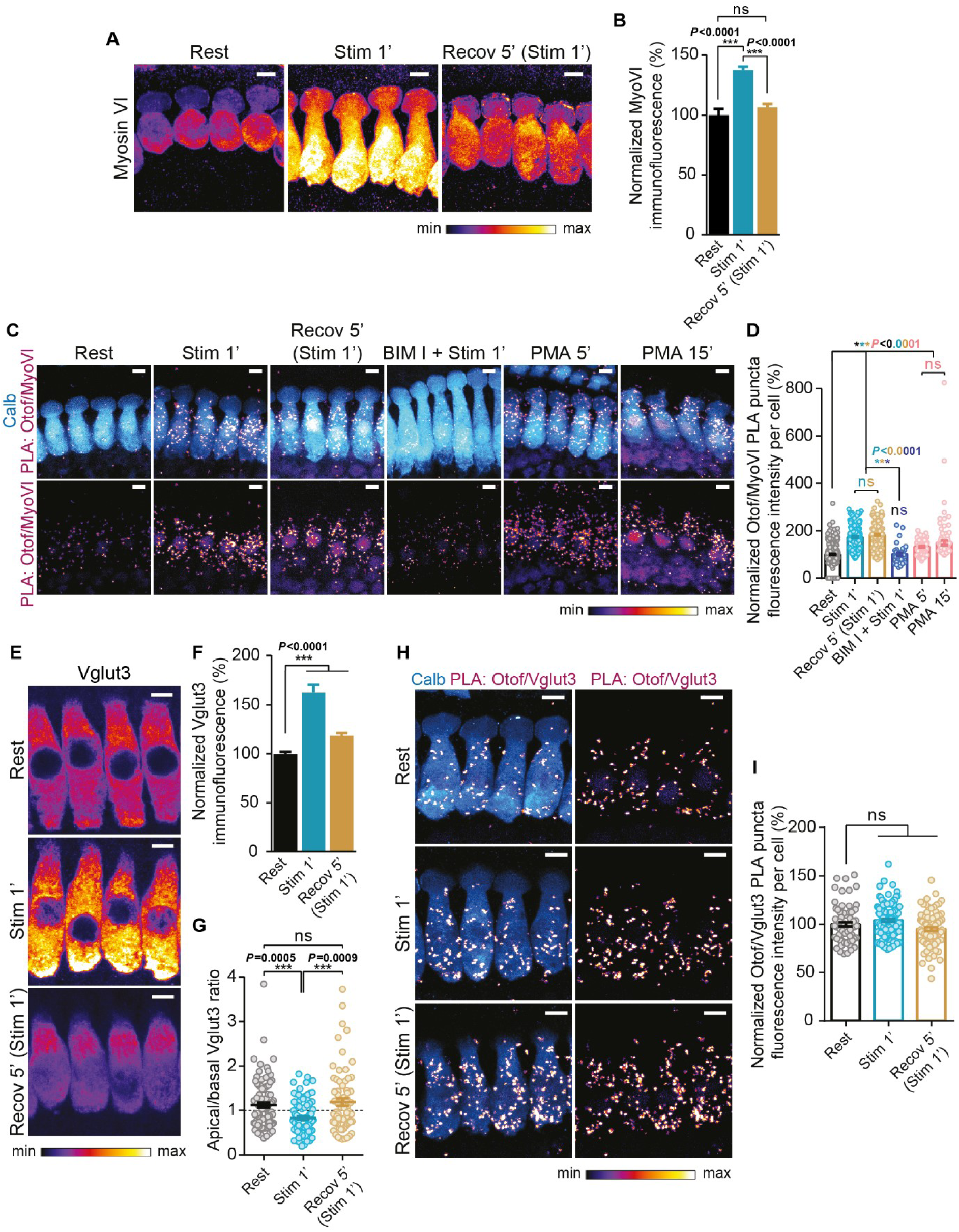
Otoferlin interacts with myosin VI, but not with Vglut3, in a PKCα-dependent manner. **A, B** Myosin VI immunofluorescence in WT P14-16 IHCs for all displayed conditions. (A) High magnification views of representative WT IHCs immunolabelled for myosin VI (intensity-coded lookup table). (B) Quantification of overall myosin VI immunofluorescence normalized to the resting condition. **C, D** PLA for otoferlin and myosin VI in WT P14-16 IHCs for all indicated conditions. (C) High magnification views of representative PLAs. (D) Average otoferlin/myosinVI PLA puncta fluorescence intensity per cell, normalized to the resting condition. See control PLAs in *Appendix Figure S2A and F*. **E-G** Vglut3 immunofluorescence in WT P14-16 IHCs for all displayed conditions. (E) High magnification views of representative WT IHCs immunolabelled for Vglut3 (intensity-coded lookup table). (F) Quantification of overall Vglut3 immunofluorescence, normalized to the resting condition. (G) Apical/basal Vglut3 immunofluorescence (above/below nuclear midline). Numbers of cells in (G) also apply to (F). **H, I** PLA for otoferlin and Vglut3 in WT P14-16 IHCs for all displayed conditions. (H) High magnification views of representative PLAs. (I) Average otoferlin/Vglut3 PLA puncta fluorescence intensity per cell, normalized to the resting condition. See control PLAs in *Appendix Figure S2A and E*. Data information: In (A, C, E, H), maximum intensity projections of confocal optical sections. Scale bars: 5 μm. In (C, H), calbindin (blue) was used as IHC marker; PLA channel is depicted with an intensity-coded lookup table with warmer colors representing higher pixel intensities. In (B, D, F-G, I), data are displayed as mean ± s.e.m.; ns *P*>0.05, \**P*≤0.05, \*\*\**P*≤0.001 (Kruskal-Wallis test followed by Dunn’s multiple comparison test); mean averages, sample size and statistical analysis are detailed in *Appendix Table S1*. In (D, G, I), individual cells are depicted with lighter colors and open symbols. Rest, resting; Stim 1’, 1-minute stimulation; Stim 5’, 5-minute stimulation; Recov 5’ (Stim 1’), 5-minute recovery after 1-minute stimulation; BIM I + Stim 1’, Incubation with BIM I prior to 1-minute stimulation; PMA 5’, 5-minute incubation with PMA; PMA 15’, 15-minute incubation with PMA. IHC, inner hair cell. Calb, calbindin. MyoVI, myosin VI. Otof, otoferlin. PLA, proximity ligation assay.

We then followed Vglut3, the vesicular glutamate transporter in IHCs, to probe for a potential regulatory role of PKC in exocytosis (Fig 6E-I). A 1-minute stimulation with high K^+^ led not only to an increase in Vglut3 immunofluorescence (Rest: 100±2%, n=89 IHCs vs. Stimulation 1’: 163±8%, n=92 IHCs; \*\*\**P*<0.0001 vs. Rest) but also to a relocation of the protein to the basal region of the IHCs (apical/basal ratio: Rest: 1.12±0.06 vs. Stimulation 1’: 0.83±0.04; \*\*\**P*<0.0001 vs. Rest) with a strong localization to the basolateral plasma membrane (Fig 6E, second panel). This likely reflects uncovering of the epitope and the transport of distal SVs to membrane-proximal sites with translocation of Vglut3 from SVs to the active zone membrane during exocytosis. After a 5-minute recovery period the levels (Recovery 5’: 118±2%, n=71 IHCs) and localization of Vglut3 (apical/basal ratio: Recovery 5’: 1.19±0.08) returned to initial values (Fig 6E-G). A PLA for otoferlin and Vglut3 was positive among conditions but no change in intensity was registered (Rest: 100±2%, n=78 IHCs vs. Stimulation 1’: 104±1%, n=146 IHCs vs. Recovery 5’: 95±2%, n=93 IHCs) (Fig 6H-I). This might be explained by the fact that both proteins are known to localize to common structures in IHCs (Strenzke *et al*, 2016) and therefore follow at least in part the same trafficking pathways without necessarily interacting.

### Otoferlin interacts with calbindin and this interaction is strongly dependent on PKCα

In most PLA experiments we used calbindin-D28k (henceforth, calbindin) as a hair cell marker and noticed a change in calbindin immunofluorescence among the different experimental conditions (Fig 7A-C). Calbindin is a member of the calmodulin superfamily of Ca^2+^-binding proteins and it was reported to function both as Ca^2+^ buffer and Ca^2+^ sensor (Berggard, 2002). A 1-minute high K^+^ stimulation led to a decrease in calbindin immunofluorescence, probably due to reduced epitope accessibility (Rest: 100±1%, n=296 IHCs vs. Stimulation 1’: 62±2%, n=174 IHCs; \*\*\**P*<0.0001) and treatment with BIM I blocked this effect (BIM I + Stimulation 1’: 99±3%, n=26 IHCs; Kruskal-Wallis test followed by Dunn’s multiple comparison test; Fig 7B). At the same time, calbindin redistributed to the base of the IHC upon stimulation (apical/basal ratio: Rest: 1.04±0.02 vs. Stimulation 1’: 0.84±0.03; \*\*\**P*<0.0001) and this effect was again blocked by BIM I (BIM I + Stimulation 1’: 1.14±0.34; ns *P*=0.8940 vs. Rest; Kruskal-Wallis test followed by Dunn’s multiple comparison test) (Fig 7C). Calbindin seems to regain its initial location after a 5-minute recovery period (apical/basal ratio: 1.07±0.05; ns *P*=0.6380 vs. Rest; Fig 7C), while immunofluorescence levels remained low as for the stimulatory condition (73±2%, n=141 IHCs; \*\*\**P*<0.0001 vs. Rest and \**P*=0.0152 vs. Stimulation 1’; Fig 7B).

**Figure 7.**
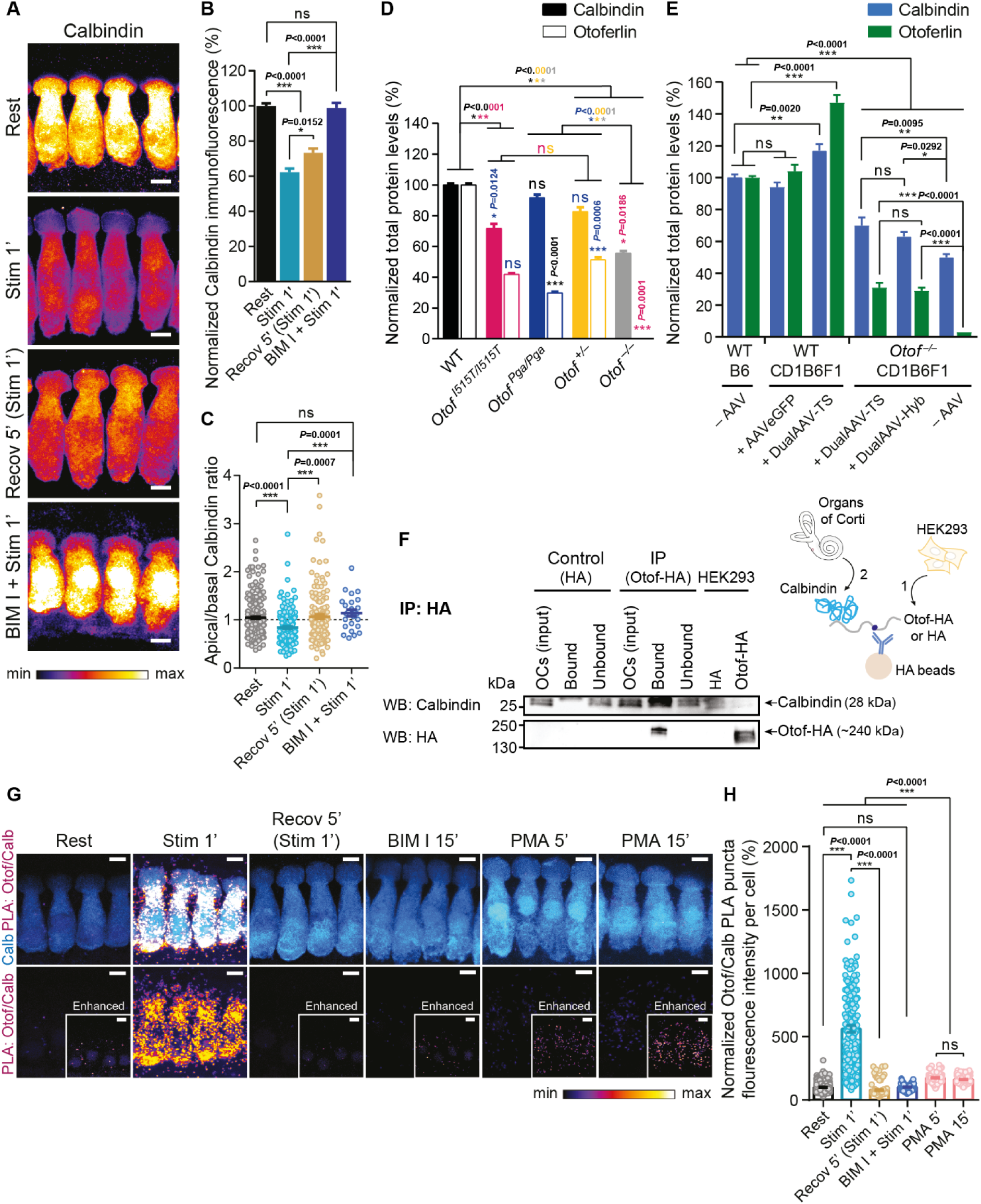
The interaction of otoferlin with calbindin is strongly dependent on PKCα. **A-C** Calbindin immunofluorescence in WT P14-16 IHCs for all indicated conditions. (A) High magnification views of representative WT IHCs immunolabelled for calbindin (intensity-coded lookup table). (B) Quantification of overall calbindin immunofluorescence. (C) Apical/basal calbindin immunofluorescence (above/below nuclear midline). Data were normalized to the resting condition. Numbers of cells in (C) also apply to (B). **D** Average calbindin and otoferlin immunofluorescence levels in otoferlin mutant and WT IHCs (P14-16). Immunofluorescence levels were normalized to WT levels for each antibody separately. See Figure EV3A for high magnification views of representative immunostainings of IHCs used for quantifications. **E** Average calbindin and otoferlin immunofluorescence levels in dual-AAV-transduced *Otof^−/−^* and WT IHCs (P23–30). Immunofluorescence levels were normalized to levels in non-transduced WT IHCs for each antibody separately. See Figure EV3B for high magnification views of representative immunostainings of IHCs used for quantifications. **F** Representative immunoblot showing results from pull-down assay from organs of Corti loaded onto anti-HA beads with previously bound otoferlin-HA expressed in HEK293 cells. Samples were probed for HA and calbindin. Right panel depicts scheme of the assay. **G, H** PLA for otoferlin and calbindin performed on WT P14-16 IHCs for all indicated conditions. (G) High magnification views of representative PLAs. (D) Average otoferlin-calbindin PLA puncta fluorescence intensity per cell for all conditions, normalized to the resting condition. See control PLAs in *Appendix Figure S2A and D*. Data information: In (A, G), maximum intensity projections of confocal optical sections. Scale bars: 5 μm. In (G), calbindin (blue) was used as IHC marker; PLA channel is depicted with an intensity-coded lookup table with warmer colors representing higher pixel intensities. In (B-E, H), data are displayed as mean ± s.e.m.; ns *P*>0.05, \**P*≤0.05, \*\**P*≤0.01, \*\*\**P*≤0.001 (Kruskal-Wallis test followed by Dunn’s multiple comparison test); mean averages, sample size and statistical analysis are detailed in *Appendix Table S1*. In (C, H), individual cells are depicted with lighter colors and open symbols. Rest, resting; Stim 1’, 1-minute stimulation; Stim 5’, 5-minute stimulation; Recov 5’ (Stim 1’), 5-minute recovery after 1-minute stimulation; BIM I + Stim 1’, Incubation with BIM I prior to 1-minute stimulation; PMA 5’, 5-minute incubation with PMA; PMA 15’, 15-minute incubation with PMA. IHC, inner hair cell. Otof, otoferlin. Calb, calbindin. PLA, proximity ligation assay.

Calbindin levels also appear to vary among several otoferlin mutants (Figs 7D and EV3A). In *Otof^−/−^* IHCs, calbindin levels were reduced to about 50% of WT levels (*Otof^−/−^*: 56±1%, n=108 IHCs vs. WT: 100±1%, n=176 IHCs; \*\*\**P*<0.0001), while IHCs of *Otof^+/–^* mice showed a reduction of about 20% (*Otof^+/–^*: 83±3%, n=99 IHCs; \*\*\**P*=0.0002; Kruskal-Wallis test followed by Dunn’s multiple comparison test; Fig 7D). For the *Otof^I515T/I515T^* mutant, carrying the temperature-sensitive p.Ile515Thr point mutation in the C_2_C domain of otoferlin (Strenzke *et al*, 2016), we found a reduction of about 25% in calbindin immunofluorescence levels when compared to WT controls (*Otof^I515T/I515T^*: 72±3%, n=83 IHCs; \*\*\**P*<0.0001), accompanied by the previously reported reduction in otoferlin levels (Fig 7D). In *Otof^Pga/Pga^* mutant IHCs (Pangršič *et al*, 2010), carrying the p.Asp1767Gly missense mutation in the C_2_F domain, there were no evident changes in calbindin levels when compared to WT IHCs (*Otof^Pga/Pga^*: 92±2%, n=76 IHCs; ns *P*=0.5900; Kruskal-Wallis test followed by Dunn’s multiple comparison test), although otoferlin levels are slightly lower in this mutant by comparison to *Otof^I515T/I515T^* IHCs (Fig 7D). In our recent study where we partially rescued hearing in *Otof^−/−^* mice by reintroducing otoferlin in IHCs via dual-AAV approaches (Al-Moyed *et al*, 2019), we quantified calbindin immunofluorescence levels alongside otoferlin levels (Figs 7E and EV3B). Reintroduction of otoferlin led to an increase in calbindin levels not only in *Otof^−/−^* IHCs (untreated *Otof^−/−^*CD1B6F1: 50±2%, n=142 IHCs; *Otof^−/−^* CD1B6F1+DualAAV-Hybrid: 63±3%, n=64 IHCs; *Otof^−/−^* CD1B6F1+DualAAV-Trans-splicing: 70±5%, n=13 IHCs), but also in wild-type IHCs (untreated WTB6: 100±2%, n=276 IHCs; WTCD1B6F1+DualAAV-Trans-splicing: 117±4%, n=62 IHCs; see Appendix Table S1 for statistics; Fig 7E).

To explore a possible interaction of otoferlin and calbindin, we repeated the otoferlin-HA pull-downs described before but this time we immunoblotted for calbindin. A strong band of ∼28 kDa in the eluate indicates a direct interaction of otoferlin and calbindin *in vitro* (Fig 7F).

We then assessed a possible interaction of otoferlin and calbindin and its potential dependency of PKCα activation in IHCs of explanted organs of Corti by a PLA (Fig 7G,H). In resting conditions, we found few PLA puncta throughout the IHCs. A 1-minute high K^+^ stimulation led to a >5-fold increase in PLA signal (Rest: 100±2%, n=327 IHCs vs. Stimulation 1’: 560±26%, n=168 IHCs; \*\*\**P*<0.0001). After a 5-minute recovery period, the PLA signal dropped to values lower than those of the resting condition (Recovery 5’: 77±5%, n=107 IHCs; \*\*\**P*<0.0001 vs. Rest). Treatment with the PKC inhibitor BIM I fully blocked the stimulation-induced increase in PLA signal (BIM I + Stimulation 1’: 101±3%, n=98 IHCs; ns *P*>0.9999 vs. Rest). Incubation with PMA led to an increase in PLA signal, though not as pronounced as for high K^+^ stimulation (PMA 5’: 175±4%, n=114 IHCs and PMA 15’: 161±3%, n=127 IHCs; \*\*\**P*<0.0001 vs. Rest and \*\*\**P*<0.0001 vs. Stimulation 1’; Kruskal-Wallis test followed by Dunn’s multiple comparison test). Thus, otoferlin and calbindin interact in IHCs in a strongly activity- and PKCα dependent manner.

A PLA between PKCα and calbindin in the same conditions (Fig EV4) resulted in an increased PLA signal after stimulation (Rest: 100±6%, n=75 IHCs vs. Stimulation 1’: 158±5%, n=94 IHCs; \*\*\**P*<0.0001), yet not as demarked as the increase observed for the PLAs between otoferlin and PKCα and between otoferlin and calbindin. This points towards an indirect interaction between calbindin and PKCα via a scaffolding protein, likely otoferlin. It is also conceivable that PKCα, otoferlin and calbindin are part of the same complex at least at some point during strong stimulation, with PKCα and calbindin binding to distinct regions of otoferlin.

## Discussion

Inner hair cells exhibit an extraordinarily high rate of synaptic vesicle turnover. Both exocytosis and endocytosis are known to be regulated by Ca^2+^ (Beutner *et al*, 2001). In this study, we found two Ca^2+^-binding proteins, PKCα and calbindin, to interact with otoferlin, thereby forming a Ca^2+^-dependent signaling complex that likely regulates different modes of endocytosis at IHC synapses.

Upon high K^+^ exposure leading to IHC depolarization, Ca^2+^ influx through voltage-gated Ca^2+^ channels triggers exocytosis, but also activates Ca^2+^-dependent kinases like PKC and CaMKII (Meese *et al*, 2017 and this study). Several proteins located next to the Ca^2+^ sources bind Ca^2+^, e.g. the proposed Ca^2+^ sensor for exocytosis at this synapse, otoferlin, as well as Ca^2+^ buffer proteins like calbindin, parvalbumin and calretinin (Pangršič *et al*, 2015). Among these, regulatory roles have so far been attributed to calbindin only (Berggard, 2002). In this study we found that PKC activation in IHCs, either pharmacologically or upon high K^+^ stimulation, triggers the interaction of PKC with otoferlin, resulting in the phosphorylation of otoferlin at S158, S775, S1169, S1224 and S1436 residues. These post-translational modifications might enable otoferlin to interact with other proteins, like myosin VI and calbindin. Pharmacological activation of PKCα without intracellular Ca^2+^ elevation also induced the interaction of otoferlin with calbindin and myosin VI, although not as effectively as by cell depolarization which triggers Ca^2+^ influx, indicating that Ca^2+^ binding to either one or both proteins strongly promotes the interaction. Since PKC inhibition before high K^+^ exposure abolished the association of otoferlin with calbindin and a direct interaction of PKCα and calbindin is rather unlikely, the phosphorylation of otoferlin by PKCα seems to be a prerequisite for the otoferlin-calbindin interaction. It is noteworthy that the increase in PLA signal for calbindin and PKC was much weaker than for the other combinations under the same stimulatory conditions, suggesting either that calbindin and PKC bind to distal parts of otoferlin, or PKC dissociates from the complex after calbindin binds to phosphorylated otoferlin.

The activation of PKC upon high K^+^ stimulation was characterized by accumulations of PKC and otoferlin in common structures near the active zones. A closer observation of the subcellular location of the interaction revealed clearly rendered fluorescent hotspots close to the synaptic ribbons. These structures were revealed to be larger than synaptic vesicles and resemble recycling endosomes described elsewhere (Kamin *et al*, 2014; Revelo *et al*, 2014; Watanabe *et al*, 2014; Jung *et al*, 2015). In an earlier study, we examined IHCs at the ultrastructural level and we found otoferlin immunogold labelling to localize to membranous compartments of >50nm diameter close to active zones, which were clearly larger than synaptic vesicles of ∼40nm. Many of the otoferlin-immunogold-labelled structures had a clathrin-coated pit at its edge, indicating these structures are most likely endosomal recycling compartments (Strenzke *et al*, 2016, Fig 7I,F,G). In addition, some otoferlin-labelled endocytic structures resembled ultrafast endocytic compartments (Strenzke *et al*, 2016, Fig 7I, F, G), which are located laterally to active zones and are about four times the size of synaptic vesicles in hippocampal synapses (Watanabe *et al*, 2013). Since ultrafast endocytosis requires a plasma membrane excess at active zones, which occurs only after strong exocytosis (Watanabe *et al*, 2013), and lower exocytosis rates rather induce clathrin mediated endocytosis (Kamin *et al*, 2014; Revelo *et al*, 2014) it is noteworthy that weak stimulation paradigms did not lead to PKCα immunofluorescence clustering in IHCs (Fig EV2A). Notably, in central nervous system synapses PKC was shown to be essential for the trafficking of synaptotagmin IX to endocytic recycling compartments (Haberman *et al*, 2005), but also seems to be involved in endocytic processes in general (Alvi *et al*, 2007). We thus propose that the structures where otoferlin and PKC interact in IHCs are most likely endocytic recycling compartments.

The nature of proteins which we found to interact with otoferlin in an activity-dependent and strongly PKCα-dependent manner supports our hypothesis that PKCα might be involved in regulating different modes of endocytosis. Upon high K^+^ IHC stimulation or treatment with a PKC activator, we observed an increase in PLA signal for the previously reported interaction of otoferlin and myosin VI (Roux *et al*, 2009; Heidrych *et al*, 2009). Myosin VI, like other myosin motors, interacts with filamentous actin (F-actin) generating the force that propels the sliding of these filaments and moves along them, thereby regulating the dynamics of the actin cytoskeleton and affecting the transport of cellular components (reviewed in Kneussel & Wagner, 2013). It was also reported that F-actin seems to control otoferlin-dependent exocytosis in auditory IHCs (Vincent *et al*, 2015), where it forms dense cage-shaped structures beneath the synaptic ribbon thereby maintaining a tight spatial organization of calcium channels at the active zones. Additionally, the authors show that F-actin colocalizes with otoferlin at the basal region of the IHC, predicting a physical association between them. Moreover, the unique myosin VI motor is involved in the early endocytic pathway, where it is required for cargo sorting (Tumbarello *et al*, 2013), so it seems plausible that both myosin VI and F-actin in association with otoferlin are involved in cellular trafficking processes in a PKC-dependent manner, which might include trafficking of endosomal compartments in IHCs.

What might be the role of calbindin in this complex? The finding that calbindin immunofluorescence is strongly reduced in *Otof^I515T/I515T^* but not in *Otof^Pga/Pga^* IHCs seems contradictory at first. Yet, a potential explanation might be that these mutations differentially impair distinct cellular processes, like vesicle replenishment (proposed for *Otof^Pga/Pga^*) and vesicle reformation from endocytic recycling compartments (ascribed to *Otof^I515T/I515T^*), and only one of these processes involves the calbindin-otoferlin interaction. In addition, a knock-out of calbindin does not affect hearing or susceptibility to noise, at least regarding threshold shifts (Airaksinen *et al*, 2000). Although a role in noise-induced synaptopathy cannot be ruled out, the short timescale of the interaction, growing weaker between 1 and 5-minute depolarizations, makes it unlikely that calbindin acts in processes that need to last from minutes to hours, such as affecting the susceptibility to noise. Similarly, triple knock-out mice of calbindin, parvalbumin and calretinin (Ca^2+^ buffer TKO) showed remarkably low impact on hearing (Pangršič *et al*, 2015). In patch clamp recordings from Ca^2+^ buffer TKO IHCs, exocytosis upon short stimuli (reflecting the fusion of the readily releasable pool of vesicles) remained wild-type-like; hence, an involvement of the Ca^2+^ buffer proteins in vesicle fusion seems unlikely. However, for longer stimuli (100-ms and 200-ms-long depolarizations to −17 mV), the change in plasma membrane capacitance (ΔC_m_) was larger in Ca^2+^ buffer TKO than in wild-type control IHCs (Pangršič *et al*, 2015, Fig 3C). Substitution of endogenous buffers with variable concentrations of the synthetic Ca^2+^ buffers EGTA or BAPTA could not accurately restore the C_m_ changes in response to fast and sustained stimuli to wild-type values, indicating that at least one of the Ca^2+^ buffer proteins might fulfill an additional function over simple Ca^2+^ buffering. At the time, the larger ΔC_m_ obtained for Ca^2+^ buffer TKO IHCs in response to longer depolarizations was presumed to reflect an increase in exocytosis, which, nonetheless, did not trigger more action potentials in postsynaptic neurons, and this apparent increase in exocytosis was then attributed to extrasynaptic vesicle fusion. Yet, 200-ms-long stimulations resulted in a ΔC_m_ of 360 fF in Ca^2+^ buffer TKO IHCs vs. 116 fF in wild-type IHCs (at 2 mM [Ca^2+^]_e_), implying that extrasynaptic exocytosis would need to occur at double the rate of synaptic exocytosis if this were the only explanation. However, corresponding amounts of extrasynaptic vesicles were never found in EM ultrastructure images, and particularly the ribbon is presumed to assist in vesicle reformation and resupply (Jung *et al*, 2015; Pangrsic & Vogl, 2018; Jean *et al*, 2018). Instead, we favor the hypothesis that ultrafast endocytosis, occurring in wild-type but absent in Ca^2+^ buffer TKO IHCs, might explain a major part of the difference in ΔC_m_ for 100-ms and 200-ms stimulations. This mode of endocytosis was first proposed by Watanabe and collaborators (Watanabe *et al*, 2013). The authors stimulated hippocampal neurons expressing channelrhodopsin with a short light pulse and fixed the tissue within few milliseconds by high-pressure quick freezing (“flash-and-freeze”). Ultrastructural analysis revealed membrane invaginations next to active zones, which were detached from the plasma membrane between 50 and 100 ms of stimulation. In C_m_ recordings, membrane invaginations do not lead to a reduction of cellular capacitance, but once the compartments become constricted and are further internalized, the plasma membrane surface area, and proportionally to it the C_m_, decrease. In the recordings of Pangrsic *et al*, ΔC_m_ from Ca^2+^ buffer TKO IHCs was larger than in wild-type IHCs, but only from 100-ms stimulations onwards (∼140 fF in Ca^2+^ buffer TKO IHCs vs. ∼60 fF in wild-type IHCs for 100-ms depolarizations at 2 mM [Ca^2+^]_e_). This could be interpreted that at least one of the Ca^2+^ buffer proteins might be required for ultrafast endocytosis.

In a follow-up study, Watanabe and collaborators found that ultrafast endocytosis depends on actin polymerization (Watanabe *et al*, 2014). When actin polymerization was inhibited with latrunculin A, the authors found a strong reduction in ultrafast endocytosis, again revealed by flash-and-freeze and EM analysis. In different studies aiming at elucidating the role of actin polymerization in IHC synaptic function (Vincent *et al*, 2015; Guillet *et al*, 2016), latrunculin A was used during C_m_ recordings of IHCs. Again, ΔC_m_ increased more in latrunculin A-treated IHCs both for whole-cell patch clamp recordings and flash photolysis of caged Ca^2+^. The authors interpreted this as facilitation of exocytosis by reduction of actin filament-based diffusion barriers and proposed a role for F-actin in controlling the diffusion rate of the synaptic vesicles to the sites of release in IHCs. However, Ca^2+^ uncaging experiments in Vincent *et al* (2015) show that there is hardly any difference in exocytic rates in the first 50 ms both in presence and absence of latrunculin A (Vincent *et al*, 2015, Fig 2D), indicating that in this experimental setting the diffusion of vesicles to the sites of release was comparable. Differences in kinetics were rather registered between 50 to 100 ms after the flash (faster increase in C_m_ for latrunculin A-treated IHCs), which is coherent with the proposed time scale for ultrafast endocytosis. We thus favor the hypothesis that the increased ΔC_m_ in presence of latrunculin A, both for step depolarizations and flash photolysis, reflects absence of ultrafast endocytosis. In Guillet *et al*, ΔC_m_ was significantly larger in presence of both actin polymerization inhibitors used, but only for 20-ms-long stimulations (∼120 fF with latrunculin A vs. ∼20 fF without, 2 mM [Ca^2+^]_e_). For longer stimuli, two actin-dependent processes might be impaired that differentially affect ΔC_m_: the impairment of ultrafast endocytosis, increasing ΔC_m_, and a reduction in vesicle replenishment, reducing ΔC_m_ in comparison to untreated cells. Both effects combined might thus have resulted in non-significantly different ΔC_m_ for 50 to 100-ms-long depolarizations.

More recently, Tertrais and collaborators blocked the fission of endocytic invaginations with the dynamin blocker dyngo-4a and observed an increase in ΔC_m_ over control values (∼50 fF vs. ∼35 fF for a train of five consecutive 20-ms depolarizations, 5 mM [Ca^2+^]_e_) (Tertrais *et al*, 2019). This would be in agreement with the assumption that IHC synapses compensate the extraordinary release rates of synaptic vesicles by ultrafast endocytosis. For endocytosis triggered by flash photolysis of caged Ca^2+^ in IHCs, time constants of 10 ms for ΔC_m_ were found, which would be even faster than reported for ultrafast endocytosis at hippocampal synapses. Although this might be a plausible scenario that would explain how this synapse compensates the extraordinarily high rates of exocytosis and compares to the effect found after 20 ms stimulation in Guillet *et al* (2016), the triggering of vesicle fusion by Ca^2+^ uncaging is a rather unphysiological strong stimulus. It increases the cellular surface by >1 pF, which would require 22 000 synaptic vesicles (of 45 aF each) per pF (Neef *et al*, 2007) to fuse with the plasma membrane and might induce endocytic mechanisms that do not typically occur in more physiological conditions. Since C_m_ recordings only reveal the sum of endocytic and exocytic events, it will be important to confirm this remarkably ultrafast kinetics of endocytosis by flash-and-freeze experiments in IHCs.

What other molecular players could be involved in ultrafast endocytosis at IHC synapses? Endophilin A was first attributed to play a role in fast bulk endocytosis, but with slower kinetics than that of ultrafast endocytosis (Watanabe & Boucrot, 2017). More recently, endophilin A and synaptojanin were found to accelerate ultrafast endocytosis at hippocampal synapses (Watanabe *et al*, 2018). In C_m_ recordings of endophilin A knock-out IHCs no apparent increase in ΔC_m_ was observed (Kroll *et al*, 2019), seemingly arguing against an involvement of endophilin A in ultrafast endocytosis at this synapse. However, chronic impairment of endocytosis and vesicle reformation will inevitably affect vesicle replenishment. Since both endophilin A and synaptojanin are known to be required not only for fission of bulk endosomes but also for clathrin uncoating of recycling vesicles, inhibition of synaptic vesicle recycling might act more strongly on synaptic function in C_m_ recordings than the slowing down of ultrafast endocytosis.

In conclusion, we showed that Ca^2+^ influx activates PKCα, which phosphorylates otoferlin, enabling it to interact with calbindin and myosin VI. We propose that the association of these proteins constitutes a molecular switch with the assembly of the otoferlin-calbindin complex being required for ultrafast endocytosis in IHCs.

## Materials and Methods

### Study approval

Animal handling and experiments complied with national animal care guidelines and were approved by the board for animal welfare of the University of Göttingen and the animal welfare office of the state of Lower Saxony, Germany.

### Animals

Wild-type C57BL/6J (B6), *Otof^I515T/I515T^* (Strenzke *et al*, 2016), *Otof^Pga/Pga^* (Pangršič *et al*, 2010) and *Otof*^−/−^ (Reisinger *et al*, 2011) mice of either gender were used. For otoferlin rescue experiments, CD1xC57BL/6N-F1 (CD1B6F1) *Otof*^−/−^ and control wild-type CD1xC57BL/6N-F1 (CD1B6F1) or wild-type C57BL/6J (B6) mice were used, as previously described (Al-Moyed *et al*, 2019). The mice were housed in social groups in individually ventilated cage (IVC) racks in a specific pathogen-free facility with free access to food and water and 12-h/12-h light/dark cycles.

### Constructs

RNA isolation and cDNA synthesis from mouse organs of Corti (OCs) were carried out as described previously (Al-Moyed *et al*, 2019). To generate eGFP-mPKCα, protein kinase C α cDNA (NM_011101.3) was amplified from the organ of Corti cDNA and subcloned into pEGFP-C2. The mCherry-P2A-mOtof-HA vector contains mCherry, a P2A peptide sequence inducing ribosome skipping (Kim *et al*, 2011), the mouse organ of Corti otoferlin coding sequence (CDS) (transcript variant 4, KX060996; NM_001313767) (Strenzke *et al*, 2016) and a hemagglutinin (HA) epitope tag (YPYDVPDYA) introduced in a region where no deleterious mutations in otoferlin were reported. To generate mCherry-P2A-HA, the HA-tagged otoferlin CDS in mCherry-P2A-mOtof-HA was replaced by an HA tag.

### Co-immunoprecipitation in HEK cells

HEK293T cells were plated at a density of 1×10^6^ cells per 10 cm dish and transfected Lipofectamine® 3000 (#L3000015, Thermo Fisher Scientific) 24h post-seeding. For GFP immunoprecipitation, cells were transfected with mCherry-P2A-mOtof-HA and eGFP-mPKCα or mCherry-P2A-mOtof-HA only (control). For HA immunoprecipitation, cells were transfected with mCherry-P2A-mOtof-HA and eGFP-mPKCα or eGFP-mPKCα only (control). Cells were harvested 72h post-transfection by washing three times in PBS (137 mM NaCl, 2.7 mM KCl and 10 mM phosphate buffer solution, pH 7.4), and lysed in NP-40 lysis buffer supplemented with protease inhibitors (10 mM Tris-HCl pH 7.5, 150 mM NaCl, 0.5 mM EDTA pH 8.0, 0.5% NP-40, protease inhibitors (#4693132001, Roche, cOmplete™, EDTA-free Protease Inhibitor Cocktail)) by pipetting extensively for 1h on ice and centrifuged at 500 x g, 4 °C for 5 min to remove cell debris. The lysates were mixed with 25 μL of anti-GFP beads slurry (GFP-Trap®_MA, #gtma-10, Chemotek) or anti-HA bead slurry (Pierce™ Anti-HA Magnetic Beads, #88836, Thermo Fisher Scientific) for GFP immunoprecipitation and HA immunoprecipitation, respectively, and incubated with gentle end-over-end mixing for 4h at RT. Beads were washed three times with dilution buffer (10 mM Tris-HCl pH 7.5, 150 mM NaCl, 0.5 mM EDTA pH 8.0) before boiling for 10 min at 70 °C. Protein complexes were resolved in 4-20% Tris-glycine gels (BIO-RAD) using PageRuler™ Plus Prestained Protein Ladder (Thermo Fisher Scientific) as a marker and transferred onto nitrocellulose membranes (GE Healthcare Life Sciences). Membranes were probed with primary antibodies mouse anti-HA (#MMS-101P, Covance, 1:1000) and mouse anti-GFP (#600-301-215, Rockland, 1:1000) followed by incubation with secondary antibody goat anti-mouse IgG-HRP (#115-035-146, Jackson ImmunoResearch, 1:2000). Immobilon Forte Western HRP substrate (#WBLUF0100, Millipore) was used for detection. Protein concentration was determined with Pierce™ BCA Protein Assay Kit (#23227, Thermo Fisher Scientific).

### Pull-down assays

HEK293T cells were plated at a density of 1×10^6^ cells per 10 cm dish and transfected 24h post-seeding with mCherry-P2A-mOtof-HA or mCherry-P2A-HA (control) using Lipofectamine® 3000 (#L3000015, Thermo Fisher Scientific). Cells were harvested 72h post-transfection by washing three times in PBS and lysed in NP-40 lysis buffer supplemented with protease inhibitors by pipetting extensively every 10 min for 1h on ice and centrifuged at 500 x g, 4 °C for 5 min to remove cell debris. Lysates were mixed with 25 μL of anti-HA bead slurry (Pierce™ Anti-HA Magnetic Beads, #88836, Thermo Fisher Scientific) incubated with gentle end-over-end mixing for 1h at 4 °C. Beads were washed three times with dilution buffer (10 mM Tris-HCl pH 7.5, 150 mM NaCl, 0.5 mM EDTA pH 8.0).

OCs from 25 mice at P8-P9 were homogenized in ice-cold sucrose buffer (320 mM sucrose, 4 mM HEPES, pH 7.4, supplemented with protease inhibitors (#4693132001, Roche, cOmplete™, EDTA-free Protease Inhibitor Cocktail)) using a glass-Teflon homogenizer, with 10 strokes at 900 r.p.m (adapted from (Huttner *et al*, 1983; Hell & Jahn, 2006; Ahmed *et al*, 2013)). The homogenate was centrifuged at 500 x g, 4 °C for 5 min to remove bone and cell debris. Homogenates (500 μg total protein) were loaded onto anti-HA beads previously immobilized with HA or otoferlin-HA proteins, and incubated with gentle end-over-end mixing overnight at 4 °C. Beads were washed three times with dilution buffer prior to elution by boiling for 10 min at 70 °C.

Protein complexes were resolved in 4-20% Tris-glycine gels (BIO-RAD) using PageRuler™ Plus Prestained Protein Ladder (Thermo Fisher Scientific) as a marker and transferred onto nitrocellulose membranes (GE Healthcare Life Sciences). Membranes were probed with primary antibodies rabbit anti-PKC alpha (#ab32376, Abcam, 1:1000), mouse anti-calbindin D-28K (#CB300, Swant, 1:1000) and mouse anti-HA (#MMS-101P, Covance, 1:1000) followed by incubation with secondary antibodies goat anti-rabbit IgG (H+L)-HRP (#111-035-144, Jackson ImmunoResearch, 1:2000), goat anti-mouse Fcγ fragment specific-HRP (#115-035-008, Jackson ImmunoResearch, 1:2000), goat anti-mouse IgG (H+L)-HRP (#115-035-146, Jackson ImmunoResearch, 1:2000), respectively. Immobilon Forte Western HRP substrate (#WBLUF0100, Millipore) was used for detection. Protein concentration was determined with Pierce™ BCA Protein Assay Kit (#23227, Thermo Fisher Scientific).

### *In vitro* phosphorylation assay and mass spectrometry analysis

HA-tagged otoferlin was overexpressed in HEK293T cells and immobilized onto anti-HA beads (Pierce™ Anti-HA Magnetic Beads, #88836, Thermo Fisher Scientific) as already described. To obtain dephosphorylated otoferlin-HA, after extensive washing in dilution buffer (10 mM Tris-HCl pH 7.5, 150 mM NaCl, 0.5 mM EDTA pH 8.0), beads were incubated with equimolar amounts of alkaline phosphatase (Calf Intestinal Alkaline Phosphatase, #18009019, Thermo Fisher Scientific) in dephosphorylation buffer (50 mM Tris-HCl pH 8.5, 0.1 mM EDTA) for 30 min at 37 °C. The reaction was stopped by incubation with phosphatase inhibitors (PhosSTOP EASYpack Roche, #4906845001, Thermo Fisher Scientific) for 15 min at 25 °C. After extensive washing in dilution buffer to remove any residual phosphatase, half the sample was set aside (dephosphorylated control sample) and the other half proceeded for incubation with PKC. The kinase assay was carried out with equimolar amounts of recombinant PKCα (Recombinant human PKC alpha protein, #ab55672, Abcam) in kinase buffer (20 mM HEPES pH 7.5, 2mM DTT, 2 mM CaCl_2_, 5mM MgCl_2_, 200 nM phorbol 12-myristate 13-acetate, 260 μM phosphatidylserine, 100 μM ATP) for 30 min at 37 °C. The reaction was terminated by adding 4X NuPAGE LDS Sample Buffer supplemented with 10% beta-mercaptoethanol to the samples and boiling at 95 °C for 10 min. Protein samples were loaded onto a 4-12% NuPAGE Novex Bis-Tris Minigels (Invitrogen). Following detection by Coomassie staining, protein bands were cut out, chopped and subjected to reduction with dithiothreitol, alkylation with iodoacetamide and finally overnight digestion with trypsin. Tryptic peptides were extracted from the gel, the solution dried in a Speedvac and kept at −20°C for further analysis (Atanassov & Urlaub, 2013).

Protein digests were analyzed on a nanoflow chromatography system (Eksigent nanoLC425) hyphenated to a hybrid triple quadrupole-TOF mass spectrometer (TripleTOF 5600+) equipped with a Nanospray III ion source (Ionspray Voltage 2400 V, Interface Heater Temperature 150°C, Sheath Gas Setting 12) and controlled by Analyst TF 1.7.1 software build 1163 (all AB Sciex). In brief, peptides were dissolved in loading buffer (2% acetonitrile, 0.1% formic acid in water), enriched on a micro pillar array trapping column (1 cm, µPac, 5 µm, PharmaFluidics) and separated on an analytical micro pillar array column (200 cm, µPac, 2.5 µm, PharmaFluidics) using a 60 min linear gradient of 5-40 % acetonitrile/0.1% formic acid (v:v) at 450 nl min-1.

Qualitative LC-MS/MS analysis was performed using a Top20 data-dependent acquisition method with an MS survey scan of *m/z* 350–1250 accumulated for 250 ms at a resolution of 30,000 full width at half maximum (FWHM). MS/MS scans of *m/z* 180–1600 were accumulated for 85 ms at a resolution of 17,500 FWHM and a precursor isolation width of 0.7 FWHM, resulting in a total cycle time of 2.0 s. Precursors above a threshold MS intensity of 125 cps with charge states 2+, 3+, and 4+ were selected for MS/MS, the dynamic exclusion time was set to 45 s. MS/MS activation was achieved by CID using nitrogen as a collision gas and the manufacturer’s default rolling collision energy settings. Two technical replicates per sample were analyzed.

Protein identification was achieved using Mascot Software 2.6 (Matrixscience). LC-MS/MS runs were searched against the UniProtKB *Mus musculus* reference proteome (revision 12-2017, 60,769 entries). The search was performed with trypsin as enzyme and iodoacetamide as cysteine blocking agent. Up to two missed tryptic cleavages, methionine oxidation and S/T/Y phosphorylation as variable modifications were allowed for. Search tolerances were set to 20 ppm for the precursor mass and 0.05 Da for fragment masses, and ESI-QUAD-TOF specified as the instrument type. Extracted Ion Chromatograms (XICs) were generated in PeakView Software version 2.1 build 11041 (AB Sciex) using 0.05 *m/z* extraction windows.

### Immunohistochemistry and Proximity ligation assay

For general immunostainings (Fig EV1) and quantification of total protein levels (Figs 4A-B, 7D-E, EV3), the apical turn of OCs from P14-16 mice was freshly dissected in phosphate buffered saline (PBS), directly fixed with 4% formaldehyde (FA) in phosphate buffered saline (PBS) for 45 min at 4 °C.

In otoferlin rescue experiments, cochleae of P23-30 mice were directly fixed with 4% FA in PBS for 45 min at 4 °C and decalcified in 0.12 M EDTA (pH 8.0) for 2-3 days before dissection of the OCs.

Chemical stimulation was performed essentially as described before (Kamin *et al*, 2014; Revelo *et al*, 2014). The apical turn of the OC from P14-16 mice was dissected in Hank’s Balanced Salt Solution without calcium HBSS (HBSS without Ca^2+^; composed of 5.36 mM KCl, 141.7 mM NaCl, 1 mM MgCl_2_, 0.5 mM MgSO_4_, 10 mM HEPES, 3.4 mM L-glutamine, and 6.9 mM D-glucose, pH 7.4) and then subjected to one of the following experimental conditions: i) Resting, 1 min in HBSS without Ca^2+^; b) Stimulation, 1 min in HBSS high K^+^ (KCl increased to 65.36 mM, NaCl reduced to 79.7 mM, and 2 mM CaCl_2_); c) Recovery, same as stimulated, followed by incubation for 5 min in HBSS with Ca^2+^ (NaCl reduced to 139.7 mM plus 2 mM CaCl_2_, with 5.36 mM KCl). To pharmacologically activate PKC, OCs were dissected in HBSS without Ca^2+^ and incubated for 1, 5 and 15 min in HBSS with Ca^2+^ supplemented with 1 μM PMA (phorbol 12-myristate 13-acetate; #ab120297, Abcam). To inhibit PKC, OCs were incubated for 15 min in HBSS with Ca^2+^ supplemented with 10 μM BIM I (Bisindolylmaleimide I, #203290, Merck) prior to stimulation with HBSS high K^+^ + BIM I. To pharmacologically inhibit both PKC and CaMKII, OCs were incubated for 15 min in HBSS with Ca^2+^ supplemented with 10 μM BIM I and 50 μM CaMKII inhibitor KN-93 (#Cay13319, Cayman Chemical) prior to stimulation with HBSS high K^+^ + BIM I + KN-93. All incubations were carried out at 37 °C and all solutions were prewarmed at 37 °C. OCs were subsequently fixed with 4% FA in PBS for 45 min at 4 °C.

Immunostainings were performed as previously described (Strenzke *et al*, 2016). The following primary antibodies were used: mouse anti-otoferlin [13A9] (#ab53233, Abcam, 1:300), rabbit anti-PKC alpha [Y124] (#ab32376, Abcam, 1:300), rabbit anti-calbindin D28k (#CB-38a, Swant, 1:300), goat anti-calbindin D28k [C-20] (#sc-7691, Santa Cruz Biotechnology, 1:100), rabbit anti-Vglut3 (#135 203, Synaptic Systems, 1:300), rabbit anti-myosin VI (KA-15) (#M5187, Sigma-Aldrich, 1:300), and goat IgG anti-CtBP2 [E-16] (#sc-5967, Santa Cruz Biotechnology, 1:100) to label the synaptic ribbons. The following secondary antibodies were used: Alexa Fluor 488-conjugated goat anti-mouse IgG (#A11001, Thermo Fisher Scientific, 1:200), Alexa Fluor 594- and Alexa Fluor 568-conjugated donkey anti-mouse IgG (#A21203, #A10037, Thermo Fisher Scientific, 1:200), Alexa Fluor 568-conjugated goat anti-rabbit IgG (#A11011, Thermo Fisher Scientific, 1:200), Alexa Fluor 488-conjugated donkey anti-rabbit IgG (#A21206, Thermo Fisher Scientific, 1:200), DyLight 405-conjugated donkey anti-goat IgG (#705-475-003, Jackson ImmunoResearch, 1:200), and MFP 488-conjugated donkey anti-goat IgG (#MFP-A1055, MoBiTec, 1:200).

Proximity ligation assay (Duolink, Sigma-Aldrich) was performed essentially as described elsewhere (Meese *et al*, 2017). The Duolink® In Situ Detection Reagents Red set was used. The following antibody combinations were used: mouse anti-otoferlin [13A9] (#ab53233, Abcam, 1:500) with rabbit anti-PKC alpha [Y124] (#ab32376, Abcam, 1:500) or rabbit anti-phosphoserine (#9332, Abcam, 1:300) or rabbit anti-calbindin D28k (#CB-38a, Swant, 1:500) or rabbit anti-Vglut3 (#135 203, Synaptic Systems, 1:500) or rabbit anti-myosin VI (KA-15) (#M5187, Sigma-Aldrich, 1:500); mouse anti-calbindin D28k (#CB300, Swant, 1:500) with rabbit anti-PKC alpha [Y124] (#ab32376, Abcam, 1:500). To visualize hair cells, primary antibody goat anti-calbindin D28K [C-20] (#sc-7691, Santa Cruz Biotechnology, 1:100) was combined with secondary antibody MFP488 donkey anti-goat IgG (#MFP-A1055, MoBiTec, 1:200), or primary antibody guinea pig anti-Vglut3 (#135 204, Synaptic Systems, 1:500) was combined with secondary antibody DyLight 405-conjugated donkey anti-guinea pig IgG (#706-475-148, Jackson ImmunoResearch, 1:100).

### Confocal microscopy and image analysis

Confocal images were acquired using a laser scanning confocal microscope Leica TCS SP5 (Leica Microsystems GmbH, Wetzlar, Germany) with a 10x air objective (0.4 NA) and a 63x glycerol-immersion objective (1.3 NA) for low and high magnification images, respectively. Exceptionally, confocal images in Fig 2 were acquired with a laser scanning confocal microscope Zeiss LSM800 with Airyscan (Carl Zeiss AG, Oberkochen, Germany) with a 63x oil-immersion objective (1.4 NA). All images from the same series were acquired with the same voltage/offset/pinhole settings and laser power.

Maximum intensity projections of optical confocal sections and single-stack images were generated using Fiji (Schindelin *et al*, 2012, https://fiji.sc/) and assembled for display in Adobe Illustrator (Adobe Systems). Color-coded 2D images were constructed in Fiji as 16-bit grayscale images to which the given color look-up table was applied. Colocalization analysis was performed using the “Coloc2” Fiji’s plugin with Costes’ autothreshold method (Costes *et al*, 2004).

Protein expression levels, immunofluorescence and PLA signals were quantified from high magnification 3D IHC images (0.6 µm z-stack step size, 2X digital zoom) in Imaris 7.6.5. Protein expression levels were quantified using a custom written Matlab (Mathworks) routine integrated into Imaris as previously described (Strenzke *et al*, 2016). PLA puncta were identified via “Spots” tool as objects with a signal above a minimum threshold. IHCs were identified by calbindin or Vglut3 fluorescence and the “Surface” tool was used to create a volume for each individual cell. Puncta per cell were obtained via the “Split Spots Into Surface Objects” Matlab XTension from Imaris, which creates a new subset of Spots that contains only the Spots that lie inside each Surface (i.e. each cell). Summed fluorescence intensities of puncta per cell were used to calculate the PLA puncta fluorescence intensity per cell and were normalized to the resting condition. For each experimental condition at least two independent experiments were performed. The same experimental settings were used for each series.

### Otoferlin rescue experiments

The viral vectors used for the otoferlin rescue experiments in Figs 7E and EV3B were designed, produced, purified, and injected through the round window membrane (RWM) into the left cochlea of P5-6 wild-type (B6 and CDB6F1) control and CD1B6F1 otoferlin knock-out mice as described in (Al-Moyed *et al*, 2019). The following virus titers were used for postnatal RWM injections: AAV2/6.eGFP (1.44 x10^10^ vg/µl), otoferlin dual-AAV2/6-TS half-vectors (1:1) (1.2 x 10^10^ vg/µl), and otoferlin dual-AAV2/6-Hyb half-vectors (1:1) (1.38 x10^10^ vg/µl). AAV2/6.eGFP was used as a control virus.

Otoferlin and calbindin protein expression levels were quantified in transduced IHCs (P23-30) and normalized to protein levels in IHCs of non-injected B6 wild-type mice as in (Al-Moyed *et al*, 2019). The otoferlin protein levels were previously reported in (Al-Moyed *et al*, 2019) and were only replotted in this study to better visualize the effect of otoferlin rescue on calbindin protein levels in otoferlin dual-AAV transduced IHCs.

### Statistical analysis

Data averages from at least two independent trials are depicted as mean ± standard error of the mean (s.e.m.). P≤0.05 value was considered significant and is denoted in figures as follows: ns P>0.05; *P≤0.05; **P≤0.01; ***P≤0.001. Statistical parameters, significance, and samples size (N, animal numbers; n, cell numbers) are reported in the figure legends. All data fitting and statistical analysis was performed using GraphPad Prism 7.03 (GraphPad Software). The D’Agostino-Pearson omnibus and the Shapiro-Wilk tests were used to test for normality. The Mann-Whitney test was used to test for statistical significance between two unpaired non-normally distributed groups. The Kruskal-Wallis test followed by the Dunn’s multiple comparison test was used to test for statistical significance in non-parametric multiple comparisons.

## Supporting information

Expandend View Figures and Appendix

## Data availability

Raw data produced in this study are available upon request.

The mass spectrometry proteomics data have been deposited to the ProteomeXchange Consortium (http://proteomecentral.proteomexchange.org) via the PRIDE partner repository (Perez-Riverol *et al*, 2019) with the dataset identifier PXD015338.

## Acknowledgments

The authors would like to thank Nina-Katrin Dankenbrink-Werder and Lisa Neuenroth for excellent technical assistance. We thank Ulrich Mueller for providing the *pachanga* mouse line. This work was supported by the University Medical Center Göttingen, the Deutsche Forschungsgemeinschaft (DFG) through the Collaborative Research Center SFB 889 (project A4 to ER and HU) and the Heisenberg Program (to ER), and the Göttingen Graduate Center for Neurosciences, Biophysics, and Molecular Biosciences (GGNB) through a stipend to APC (DFG Grant GSC 226/4).

## Author Contributions

APC and ER conceived the study. APC designed and cloned DNA constructs, carried out Co-IP, pull-down and *in vitro* phosphorylation assays, performed immunohistochemistry and proximity ligation assays, acquired and analyzed confocal microscopy images. HAM performed otoferlin rescue experiments, corresponding immunohistochemistry and confocal microscopy image acquisition and analysis. CL designed and evaluated mass spectrometry experiments. APC, HAM and CL analyzed data and prepared figures. APC and ER wrote the manuscript with input from all authors. All authors revised the manuscript. ER and HU acquired funding.

## Expanded View Figure Legends

**Figure EV1. PKCα is expressed in the organ of Corti, in IHCs and OHCs.**

**A, B** WT organ of Corti (P15) immunolabeled for otoferlin and PKCα. (A) Low magnification views. (B) High magnification views of IHCs from the organ of Corti shown in (A). PKCα channel is depicted separately with an intensity-coded lookup table with warmer colors representing higher pixel intensities. IHCs: inner hair cells, OHCs: outer hair cells.
**C, D** (C) Fluorescence intensity line profile through the longitudinal axis at the mid-region of a representative IHC, from apex to base (five optical sections). (D) Representative IHC used for the fluorescence intensity line profile. PKCα and otoferlin channels depicted separately (intensity-coded lookup table).
Data information: In (A-B), maximum intensity projections of confocal optical sections. (D) Single optical section. Scale bars: 100 μm (A), 5 μm (B), 2 μm (D).

**Figure EV2. PKCα redistributes upon strong stimulation and treatment with PMA.**

**A** High magnification views of representative WT P15-16 IHCs immunolabeled for PKCα, after mild (25 mM KCl) and strong (65 mM KCl) stimulations for 1 minute. PKCα staining is depicted with an intensity-coded lookup table with warmer colors representing higher pixel intensities.
**B** High magnification views of representative WT P15-16 IHCs immunolabeled for PKCα and otoferlin, at rest (Rest), and after treatment with the PKC activator PMA for 1 (PMA 1’), 5 (PMA 5’) and 15 (PMA 15’) minutes. Individual otoferlin and PKCα channels are depicted separately with an intensity-coded lookup table with warmer colors representing higher pixel intensities.
**C** Fluorescence intensity line profile through the longitudinal axis at the mid-region of representative IHCs labelled in (B), from apex to base (five optical sections).
Data information: In (A-B), maximum intensity projections of confocal optical sections. Scale bars: 5 μm.

**Figure EV3. Calbindin and otoferlin immunofluorescence in different otoferlin mutants and in dual-AAV-transduced *Otof^−/−^* and WT IHCs.**

**A** High magnification views of representative WT, *Otof^I515T/I515T^*, *Otof^Pga/Pga^*, *Otof^+/–^* and *Otof^−/−^* P14-16 IHCs immunolabeled for calbindin and otoferlin, used for quantification of calbindin levels in Figure 7D. Individual calbindin and otoferlin channels are depicted separately with an intensity-coded lookup table with warmer colors representing higher pixel intensities.
**B** High magnification views of dual-AAV-TS (P26) and dual-AAV-Hyb (P26) transduced CD1B6F1-*Otof*^−/−^ IHCs compared to AAV2/6.eGFP transduced WT CD1B6F1 (P28) and non-injected WT B6 (P27) IHCs, used for quantification of calbindin levels in Figure 7E. Successful virus transduction was monitored via eGFP immunofluorescence (green). Organs of Corti were immunolabeled against otoferlin (magenta) and calbindin (blue). Individual eGFP, otoferlin, and calbindin channels are depicted separately with an intensity-coded lookup table with warmer colors representing higher pixel intensities.
Data information: In (A-B), maximum intensity projections of optical confocal sections. Scale bars: 5 μm (A), 10 μm (B). IHC, inner hair cell. Calb, calbindin.

**Figure EV4. Weak PLA signal between PKCα and calbindin points toward an indirect interaction of the two proteins via scaffolding proteins.**

**A-B** PLA for PKCα and calbindin performed on WT P15-16 IHCs for the indicated conditions. (A) High magnification views of representative PLAs. (B) Average PKCα/calbindin PLA puncta fluorescence intensity per cell, normalized to the resting condition. Individual cells are depicted with lighter colors and open symbols. **C** High magnification views of representative control PLA performed with calbindin primary antibody only. See control PLA with PKCα antibody only in *Appendix Figure S2C*.
Data information: In (A, C), maximum intensity projections of confocal optical sections. Scale bars: 5 μm. Vglut3 (blue) was used as IHC marker. The PLA channel is depicted with an intensity-coded lookup table with warmer colors representing higher pixel intensities. Rest, resting; Stim 1’, 1-minute stimulation; Recov 5’ (Stim 1’), 5-minute recovery after 1-minute stimulation. Calb, calbindin.

