## Supplementary material for "PKCα-dependent interaction of otoferlin and calbindin: evidence for regulation of endocytosis in inner hair cells": Expandend View Figures and Appendix

**Figure EV1**

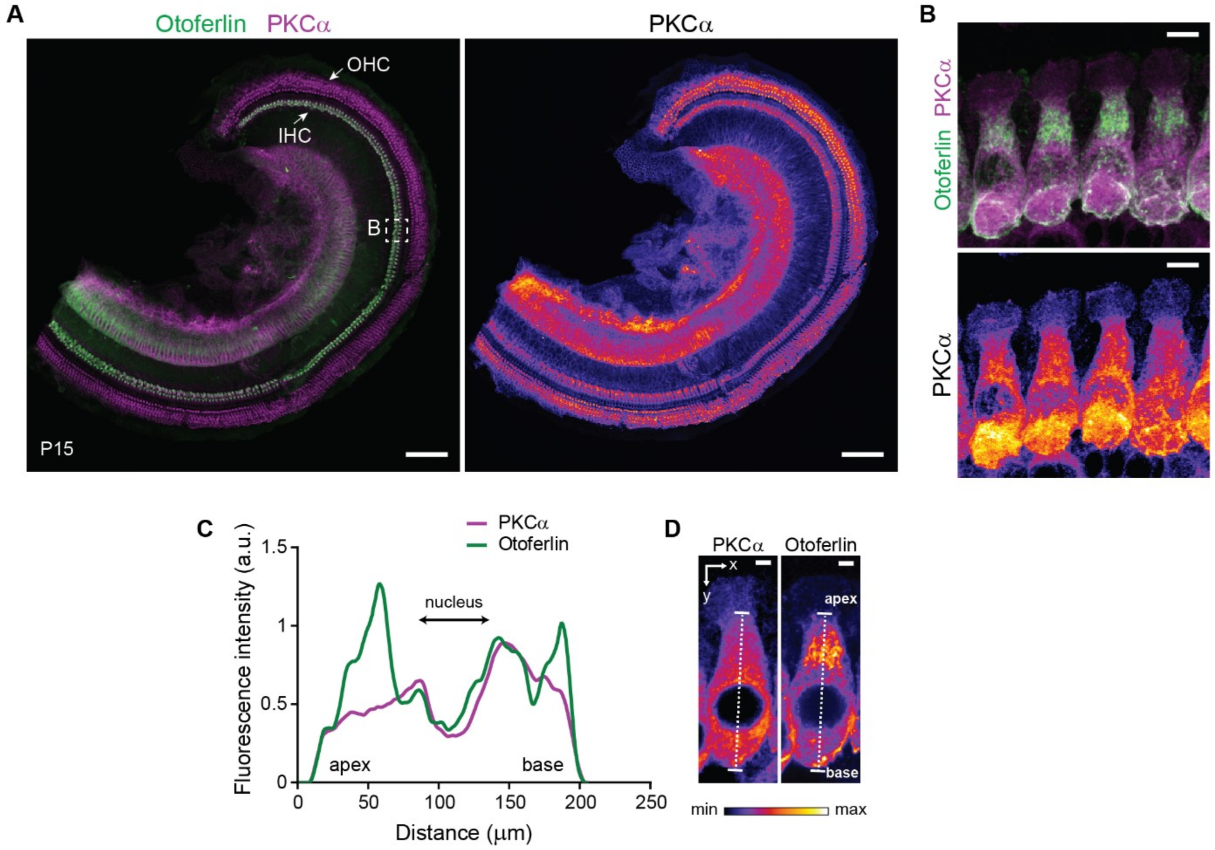
A
Otoferlin PKCα
PKCα
OHC
IHC
B
P15
B
Otoferlin PKCα
PKCα
C
PKCα
Otoferlin
Fluorescence intensity (a.u.)
1.5
1
0.5
0
nucleus
apex
base
0
50
100
150
200
250
Distance (μm)
D
PKCα
Otoferlin
x
y
apex
base
min
max

**Figure EV2**

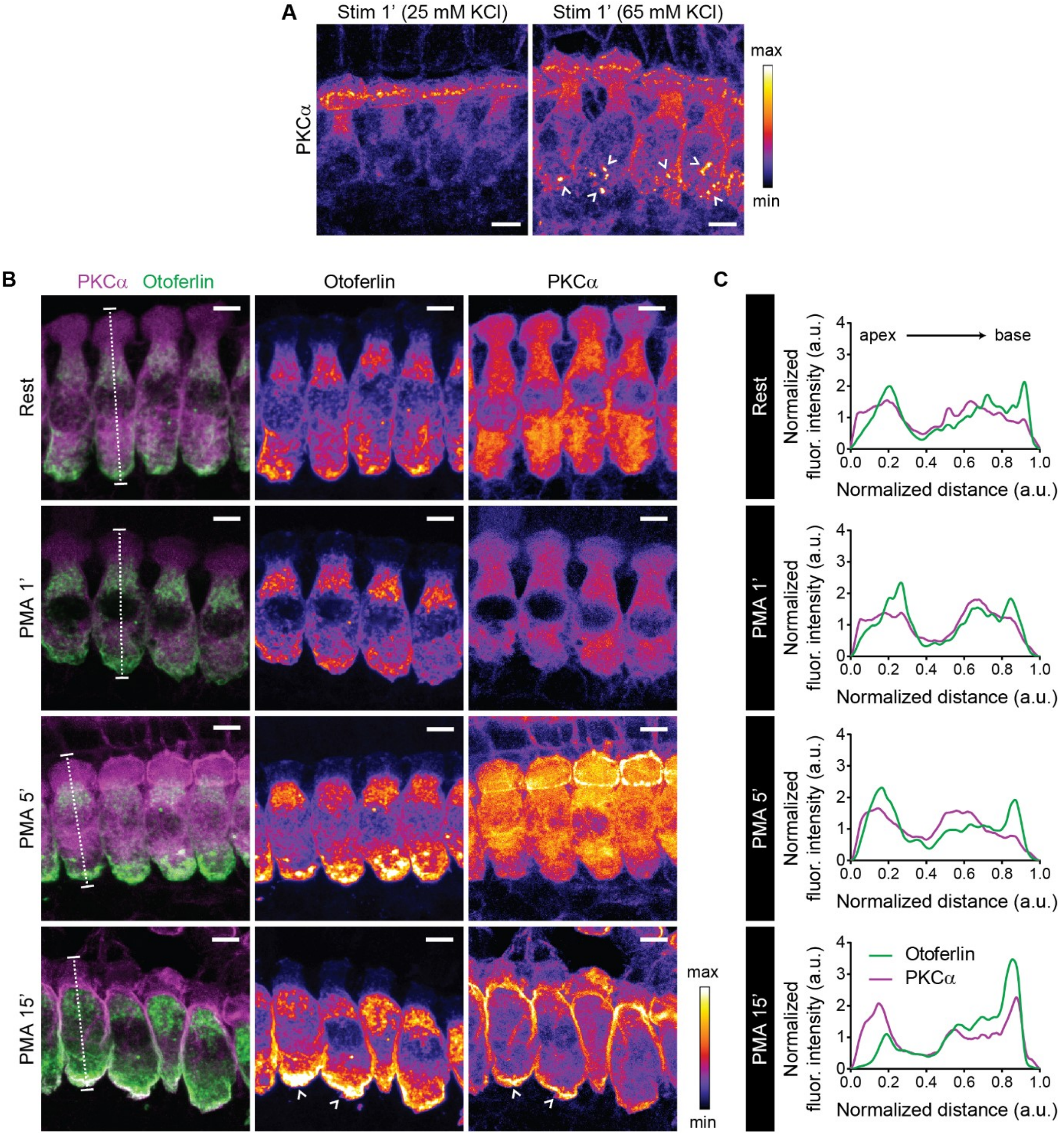
A
Stim 1' (25 mM KCl)
Stim 1' (65 mM KCl)
PKCα
max
min
B
PKCα Otoferlin
Otoferlin
PKCα
Rest
PMA 1'
PMA 5'
PMA 15'
C
Normalized fluor. intensity (a.u.)
Normalized distance (a.u.)
apex
base
Otoferlin
PKCα

**Figure EV3**

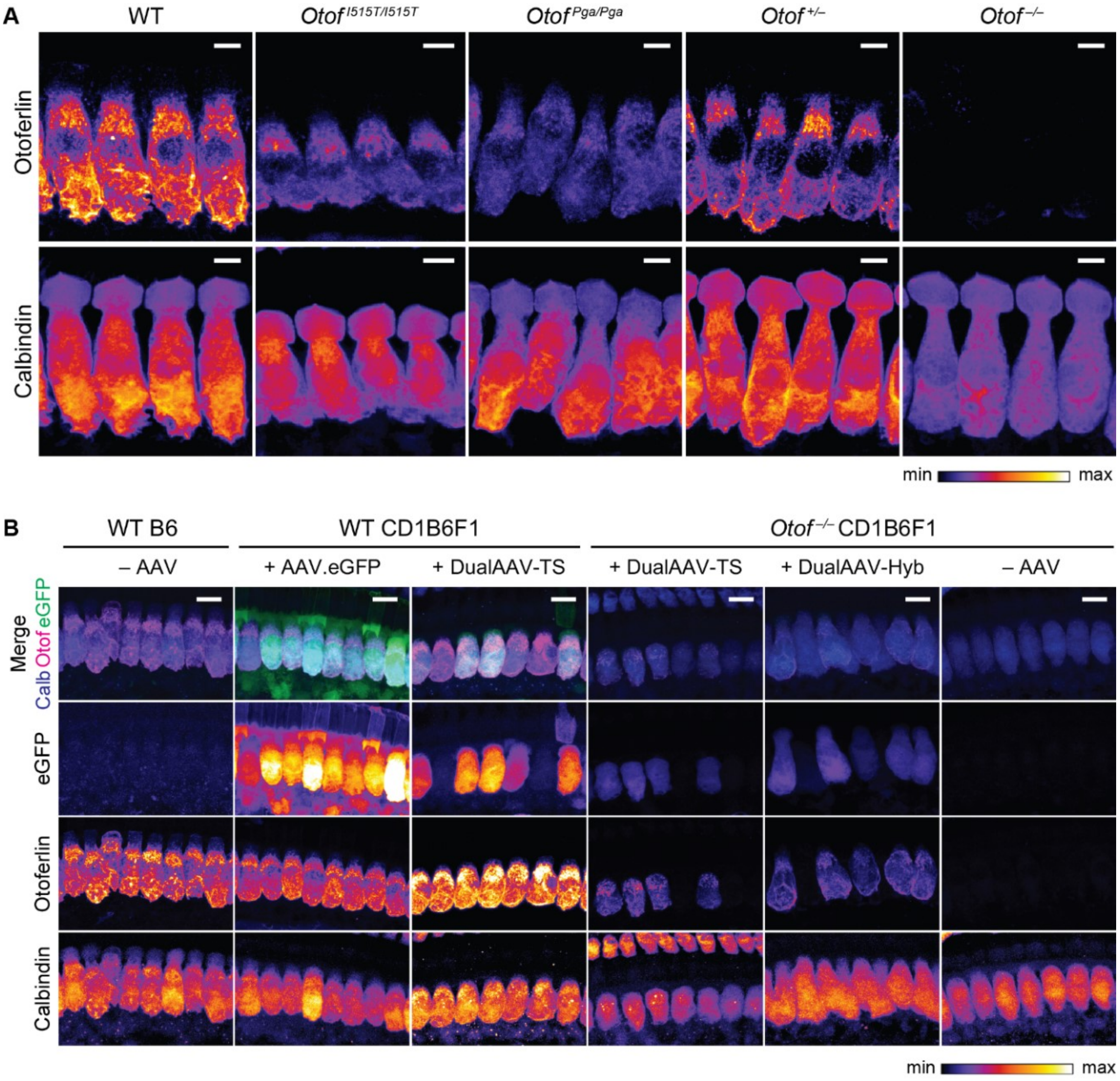
A
WT
Otof I515T/I515T
Otof Pga/Pga
Otof +/–
Otof –/–
Otoferlin
Calbindin
min
max
B
WT B6
WT CD1B6F1
Otof –/– CD1B6F1
– AAV
+ AAV.eGFP
+ DualAAV-TS
+ DualAAV-TS
+ DualAAV-Hyb
– AAV
Merge
Calb Otof eGFP
eGFP
Otoferlin
Calbindin
min
max

**Figure EV4**

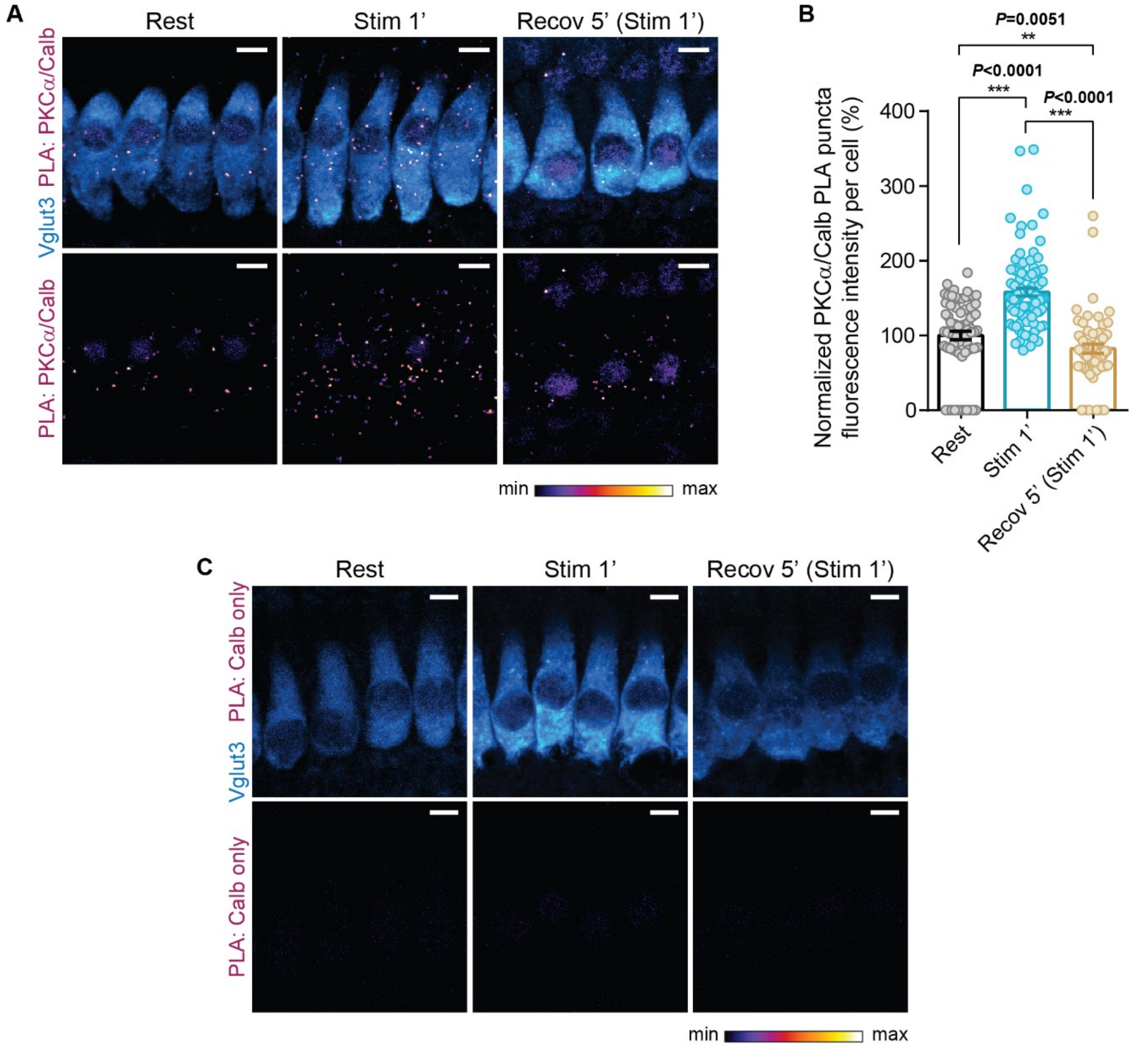
A
Rest
Stim 1'
Recov 5' (Stim 1')
Vglut3 PLA: PKCα/Calb
PLA: PKCα/Calb
min
max
B
P=0.0051
**
P<0.0001
***
P<0.0001
***
Normalized PKCα/Calb PLA puncta fluorescence intensity per cell (%)
400
300
200
100
0
Rest
Stim 1'
Recov 5' (Stim 1')
C
Rest
Stim 1'
Recov 5' (Stim 1')
Vglut3 PLA: Calb only
PLA: Calb only
min
max

### Appendix

### for

### PKCα-dependent interaction of otoferlin and calbindin: evidence for regulation of endocytosis at auditory hair cell synapses

Andreia P. Cepeda[1,2,3], Hanan Al-Moyed[1,3], Christof Lenz[4,5], Henning Urlaub[4,5], Ellen Reisinger[1,2,6*]

#### Table of contents

**Appendix Figure S1.** Validation of the proximity ligation assay in mice organs of Corti.

**Appendix Figure S2.** Negative controls for the proximity ligation assays.

**Appendix Figure S3.** MS/MS spectrum of *m/z* $632.629^{3+}$ at 38.63 min, DSQETDGLLPGSRP$^{158}$pSTR (otoferlin variant 1, NP_001093865.1).

**Appendix Figure S4.** MS/MS spectrum of *m/z* $449.209^{3+}$ at 42.93 min, FL$^{790}$pSLSDKDQGR (otoferlin variant 1, NP_001093865.1).

**Appendix Figure S5.** MS/MS spectrum of *m/z* $485.245^{3+}$ at 32.04 min, GVQS$^{1184}$pSLIHNYKK (otoferlin variant 1, NP_001093865.1).

**Appendix Figure S6.** MS/MS spectrum of *m/z* $542.614^{3+}$ at 38.08 min, YTLVGSHAVS$^{1239}$pSLRR (otoferlin variant 1, NP_001093865.1).

**Appendix Figure S7.** MS/MS spectrum of *m/z* $591.280^{2+}$ at 37.87 min, FKG$^{1451}$pSLCVYK (otoferlin variant 1, NP_001093865.1).

**Appendix Figure S8.** Total Ion Chromatograms (TICs) of otoferlin in-gel tryptic digests analyzed by LC-MS/MS

**Appendix Figure S9.** Extracted Ion Chromatograms (XICs) of otoferlin-derived phosphopeptides.

**Appendix Figure S10.** Sequence alignment of phosphorylated sites in otoferlin variants 1 (NP_001093865.1) and 4 (NP_001300696.1).

**Appendix Figure S11.** Sequence alignment of phosphorylated sites in otoferlin from different species.

**Appendix Figure S12.** PKC is predicted to phosphorylate otoferlin.

**Appendix Table S1.** Mean averages, sample size and statistical analysis.

**Appendix Table S2.** Prediction of PKC phosphorylation sites in otoferlin.

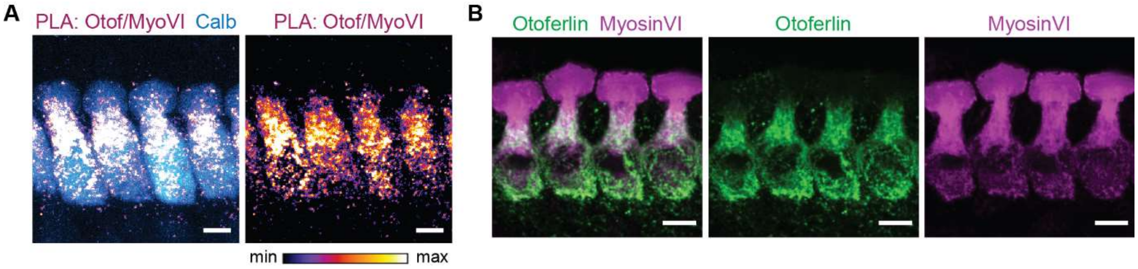

**Appendix Figure S1. Validation of the proximity ligation assay in mice organs of Corti.**

**A** High magnification views of a PLA assay for otoferlin and myosin VI performed on WT B6 P14 IHCs. Calbindin (blue) was used as IHC marker. PLA channel is depicted with an intensity-coded lookup table with warmer colors representing higher pixel intensities.

**B** High magnification views of WT B6 P14 IHCs immunolabeled for otoferlin and myosin VI with the antibodies used for the PLA shown in (A).

Data information: In (A-B), maximum intensity projections of confocal optical sections. Scale bars: 5 μm. PLA, proximity ligation assay. IHC, inner hair cell. Otof, otoferlin. MyoVI, myosin VI. Calb, calbindin.

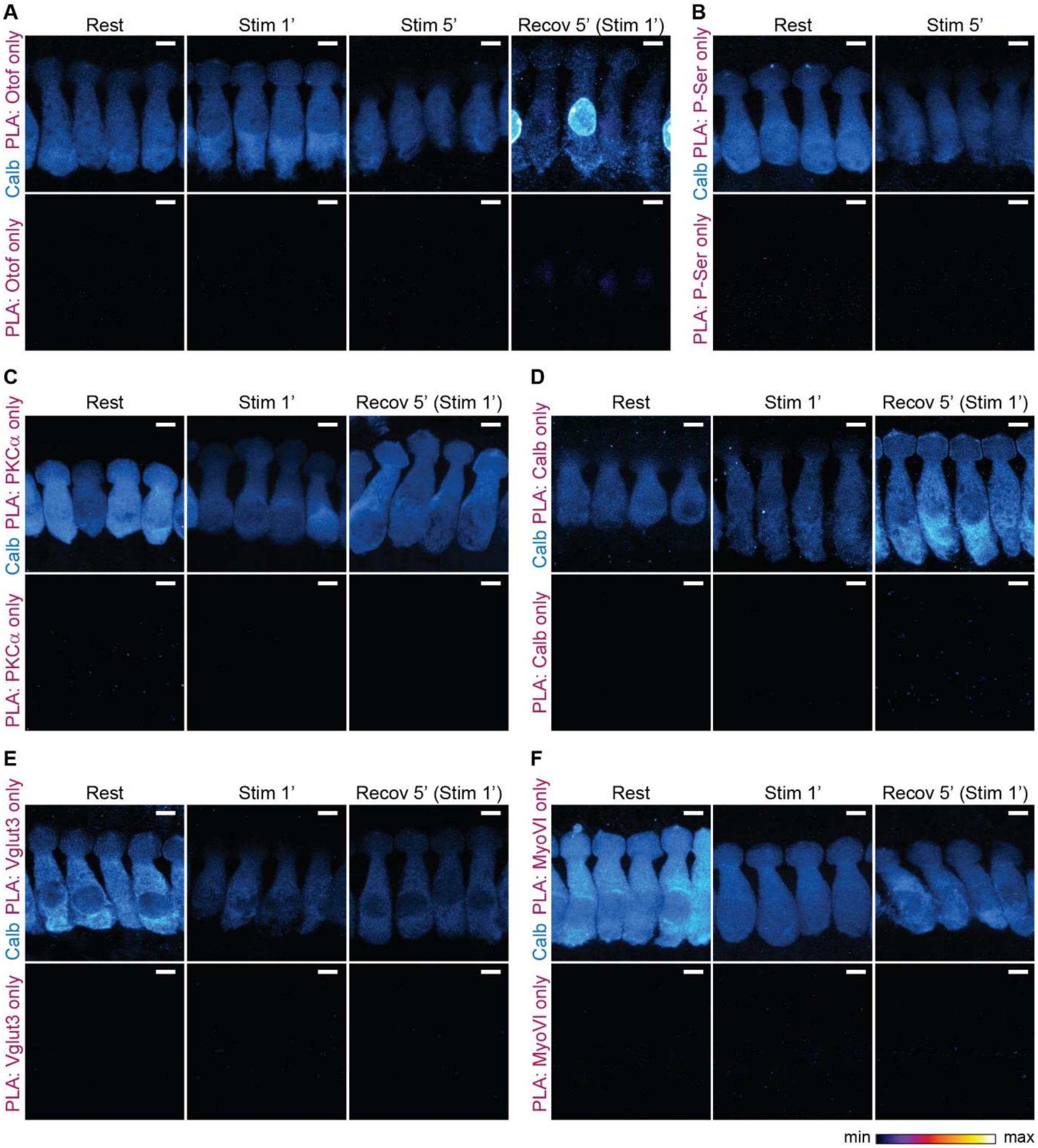

**Appendix Figure S2. Negative controls for the proximity ligation assays.**

**A-F** High magnification views of representative control PLAs performed with only one of the primary antibodies and done in parallel to the PLAs presented in this study: anti-otoferlin (A), anti-phosphoserine (B), PKC$\alpha$ (C), calbindin (D), Vglut3 (E), myosin VI (F). Calbindin (blue) was used as IHC marker. The PLA channel is depicted with an intensity-coded lookup table (fire) with warmer colors representing higher pixel intensities. PLAs were performed for the conditions where the strongest PLA signal was registered in all different PLA combinations.

Data information: In (A-F), maximum intensity projections of confocal optical sections. Scale bars: 5 $\mu$m. Rest, resting; Stim 1’, 1-minute stimulation; Stim 5’, 5-minute stimulation; Recov 5’ (Stim 1’), 5-minute recovery after 1-minute stimulation. PLA, proximity ligation assay. Calb, calbindin. Otof, otoferlin. P-Ser, phosphoserine.

### Annotated MS/MS Spectra of Detected Phosphosites

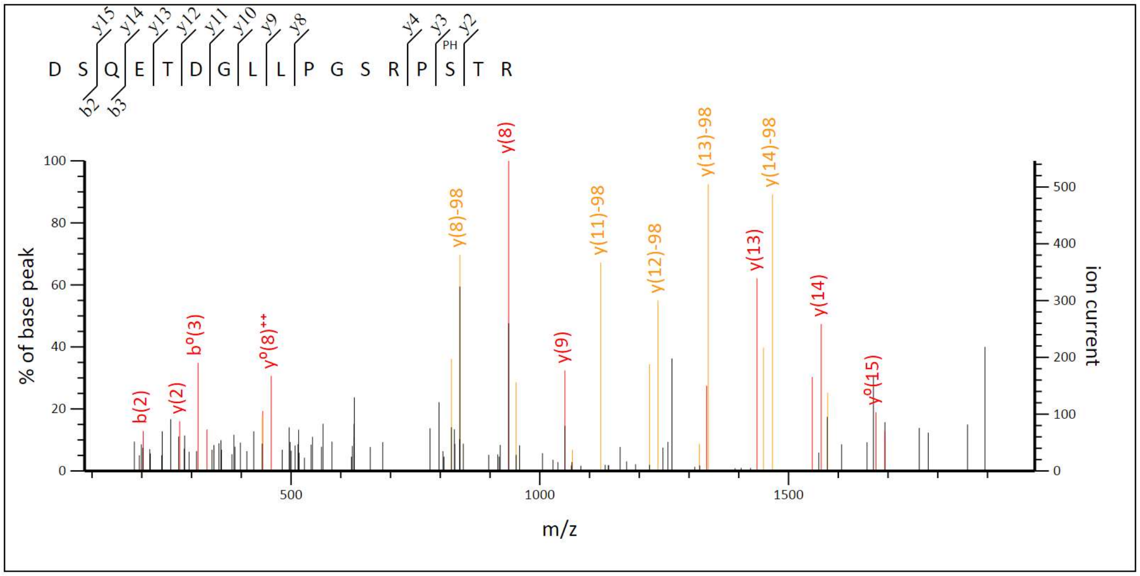

**Appendix Figure S3.** MS/MS spectrum of *m/z* $632.629^{3+}$ at 38.63 min, DSQETDGLLPGSRP$^{158}$pSTR (otoferlin variant 1, NP_001093865.1).

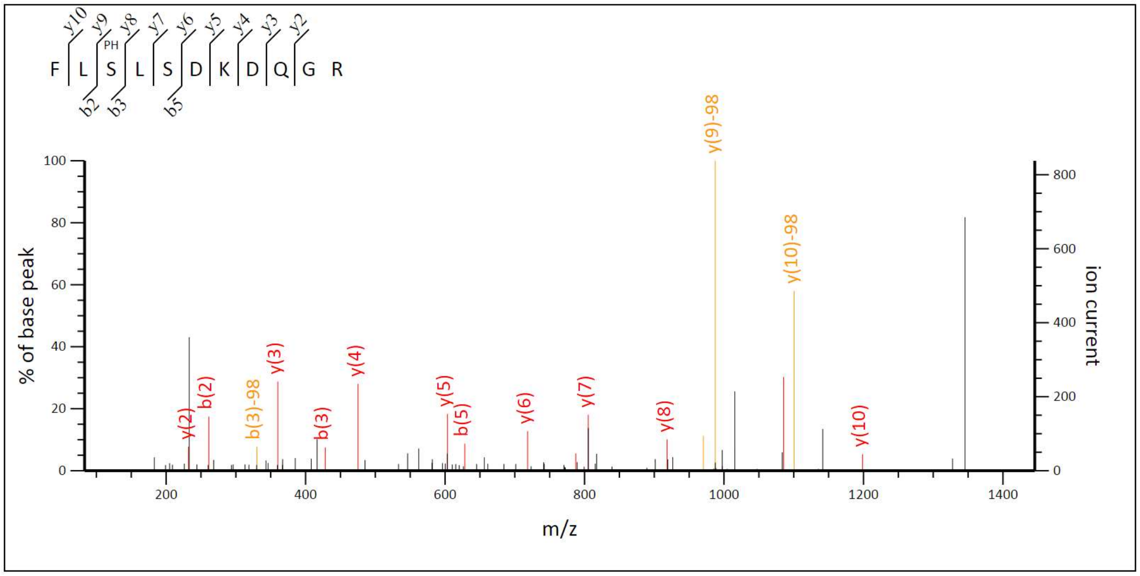

**Appendix Figure S4.** MS/MS spectrum of *m/z* $449.209^{3+}$ at 42.93 min, FL$^{790}$pSLSDKDQGR (otoferlin variant 1, NP_001093865.1).

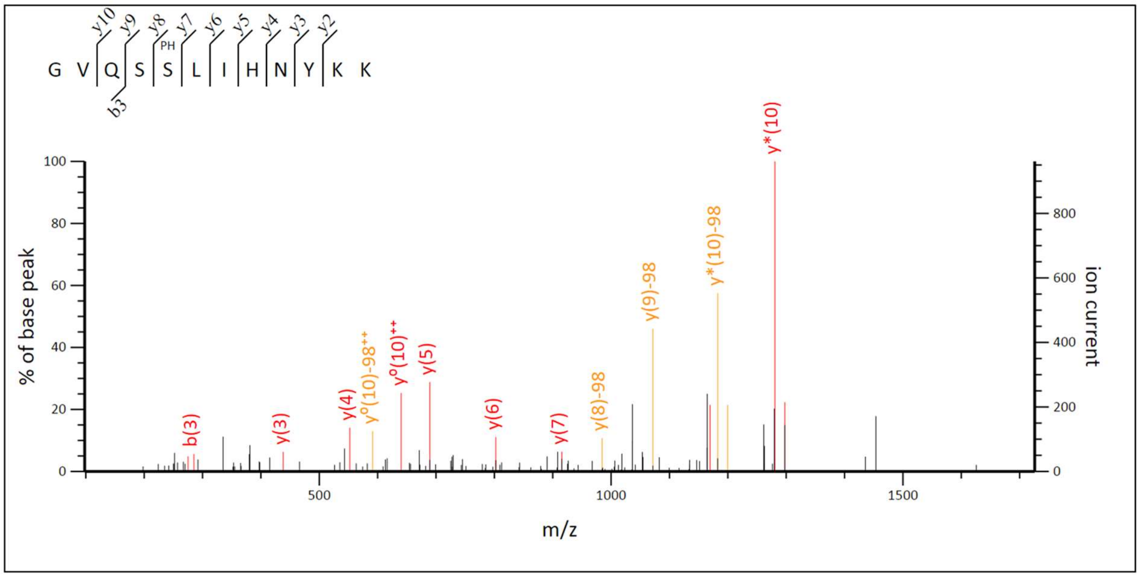

**Appendix Figure S5.** MS/MS spectrum of *m/z* $485.245^{3+}$ at 32.04 min, GVQS[1184]pSLIHNYKK (otoferlin variant 1, NP_001093865.1).

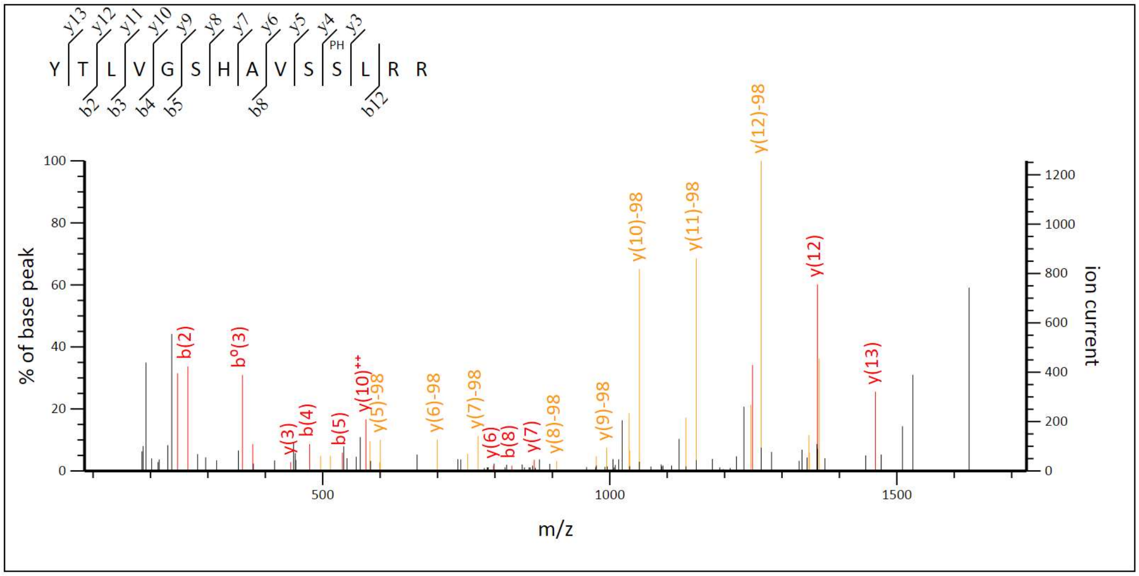

**Appendix Figure S6.** MS/MS spectrum of *m/z* $542.614^{3+}$ at 38.08 min, YTLVGSHAVS[1239]pSLRR (otoferlin variant 1, NP_001093865.1).

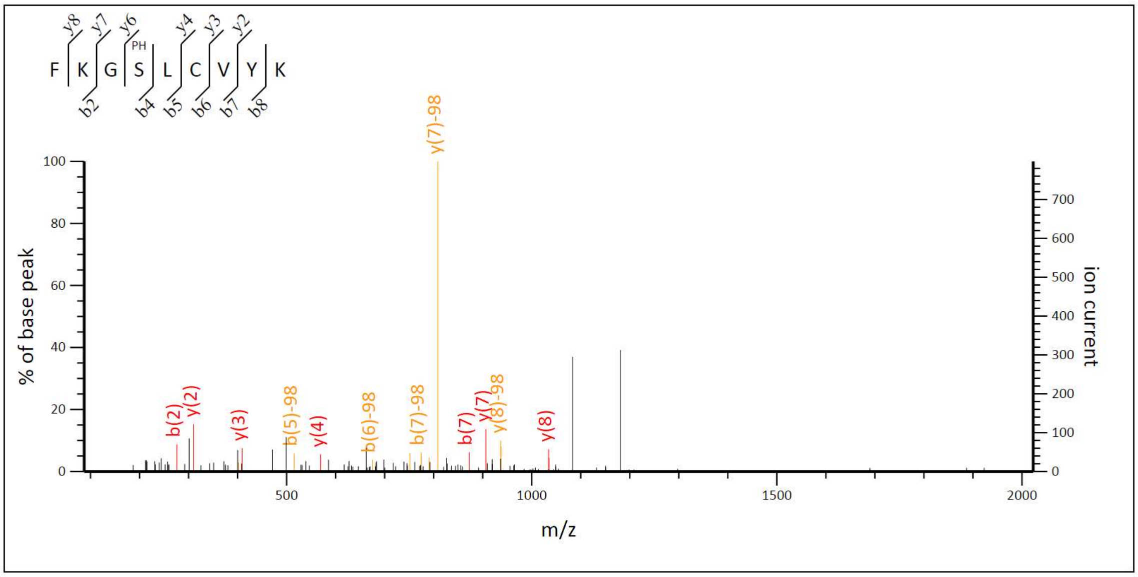

**Appendix Figure S7.** MS/MS spectrum of *m/z* 591.280$^{2+}$ at 37.87 min, FKG$^{1451}$pSLCVYK (otoferlin variant 1, NP_001093865.1).

### LC-MS/MS profiling of phosphopeptides

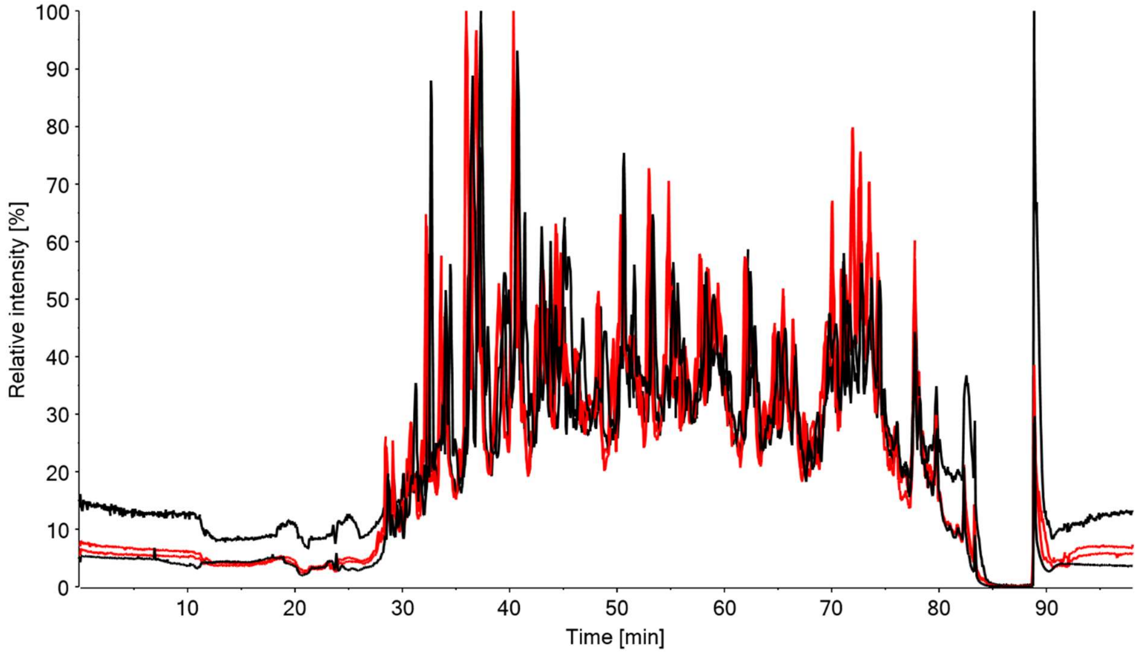

**Appendix Figure S8. Total Ion Chromatograms (TICs) of otoferlin in-gel tryptic digests analyzed by LC-MS/MS.** Two replicates each of phosphatase-treated (black) and phosphatase-treated/PKC-incubated (red) samples were analyzed. XIC overlays demonstrate excellent reproducibility.

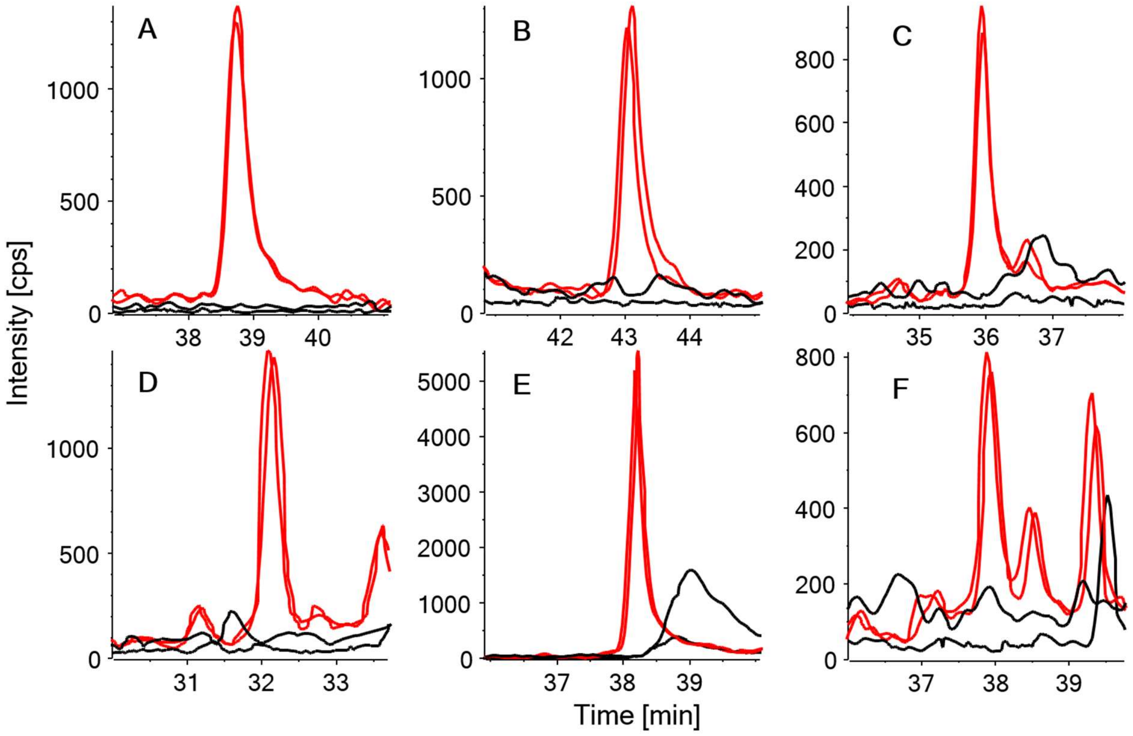

**Appendix Figure S9. Extracted Ion Chromatograms (XICs) of otoferlin-derived phosphopeptides.** Two replicates each of phosphatase-treated (black) and phosphatase-treated/PKC-incubated (red) samples were analyzed.

**A** *m/z* 632.629$^{3+}$ DSQETDGLLPGSRP$^{158}$pSTR

**B** *m/z* 449.209$^{3+}$ FL$^{790}$pSLSDKDQGR

**C** *m/z* 663.322$^{2+}$ GVQS$^{1184}$pSLIHNYK

**D** *m/z* 485.245$^{3+}$ GVQS$^{1184}$pSLIHNYKK

**E** *m/z* 542.614$^{3+}$ YTLVGSHAVS$^{1239}$pSLRR

**F** *m/z* 591.280$^{2+}$ FKG$^{1451}$pSLCVYK

Background signal in the phosphatase-treated samples (black) indicates that phosphorylation was achieved by PKC incubation (red).

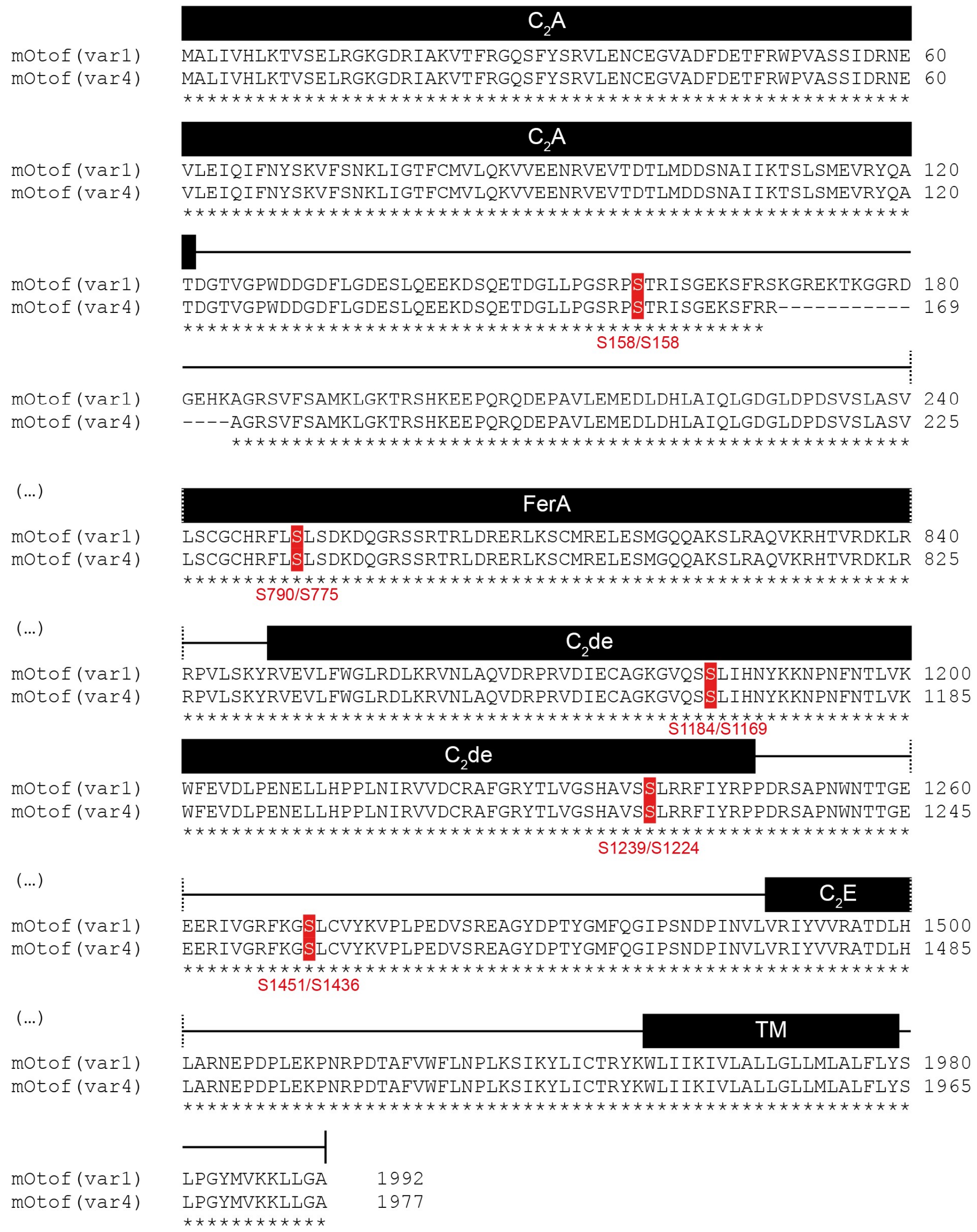

**Appendix Figure S10. Sequence alignment of phosphorylated sites in otoferlin variants 1 (NP_001093865.1) and 4 (NP_001300696.1).** Phosphosites (red) identified at positions S158, S790, S1184, S1239, S1451 in variant 1 correspond to S158, S775, S1169, S1224, S1436 in variant 4, respectively. Alignment was performed using CLUSTAL Omega (1.2.4), EMBL-EBI.

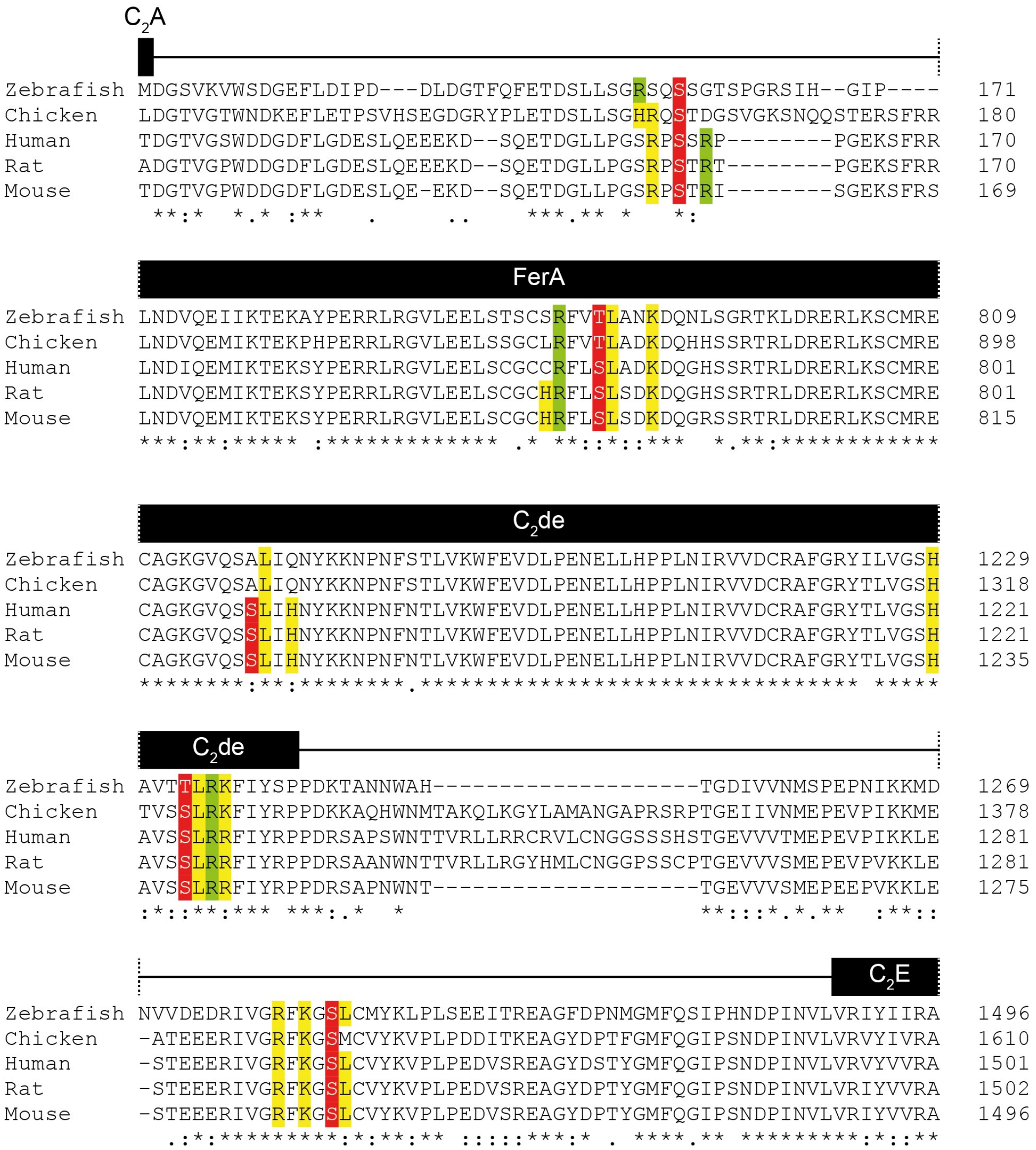

**Appendix Figure S11. Sequence alignment of phosphorylated sites in otoferlin from different species.** Identified phosphorylation sites in mouse otoferlin (NP_001093865.1, variant 1) were aligned with human (NP_919224.1), rat (NP_001263649.1), chicken (XP_015140684.1) and zebrafish (NP_001025283.1) otoferlin sequences (displayed in red). PKC consensus motif (R-X-X-S/T-X-R-X) with with respective arginines (R) indicated in green; X indicates any amino acid. Hydrophobic leucine residues (L) at the +1 position shown to be favored by PKC and reported variations to the consensus motif with basic amino acids (R, H, K) at positions −6, −4, −2, +2, +3, and +4 (Nishikawa *et al*, 1997) are depicted in yellow. Alignment was performed using CLUSTAL Omega (1.2.4), EMBL-EBI.

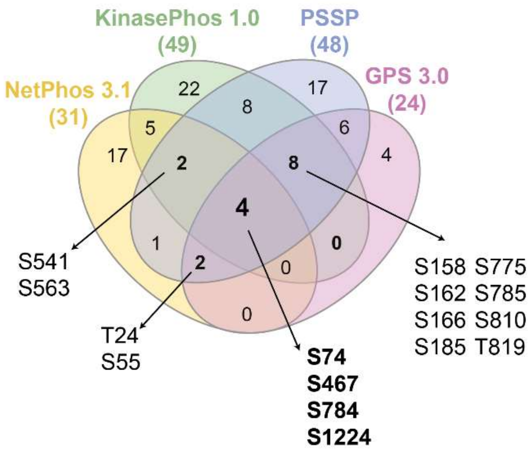

**Appendix Figure S12. PKC is predicted to phosphorylate otoferlin.** Analysis of putative PKC phosphorylation sites in otoferlin (mouse, isoform 4, NP_001300696.1) using four different prediction tools (see *Appendix Table S2* for detailed analysis). A comparative analysis between tools is represented as Venn Diagram. Numbers in parenthesis refer to total number of sites for each tool. Common sites to all tools are displayed in bold. Common sites found in at least three of the tools are also depicted.

#### Appendix Table S1. Mean averages, sample size and statistical analysis.

Data information: s.e.m., standard error of the mean; N, number of animals; n, number of cells.

**Figure 1D.**

| Genotype/Condition | Mean ± s.e.m. | | N | n |
|---|---|---|---|---|
| | Otoferlin | PKCα | | |
| Rest | 1.03 ± 0.02 | 1.05 ± 0.03 | 6 | 150 |
| Stim 1' | 0.89 ± 0.01 | 0.76 ± 0.02 | 7 | 179 |
| Stim 5' | 0.94 ± 0.06 | 0.85 ± 0.06 | 1 | 37 |
| Recov 5' (Stim 1') | 1.60 ± 0.04 | 1.32 ± 0.04 | 6 | 195 |

| Compared group | Statistical significance | *P*-value | Statistical test |
|---|---|---|---|
| ***Apical/basal Otoferlin ratio:*** | | | |
| Rest vs. Stim 1' | *** | 0.0004 | Kruskal-Wallis test followed by Dunn's multiple comparison test |
| Rest vs. Stim 5' | ns | 0.0658 | |
| Rest vs. Recov 5' (Stim 1') | *** | < 0.0001 | |
| Stim 1' vs. Stim 5' | ns | > 0.9999 | |
| Stim 1' vs. Recov 5' (Stim 1') | *** | < 0.0001 | |
| Stim 5' vs. Recov 5' (Stim 1') | *** | < 0.0001 | |
| ***Apical/basal PKCα ratio:*** | | | |
| Rest vs. Stim 1' | *** | < 0.0001 | Kruskal-Wallis test followed by Dunn's multiple comparison test |
| Rest vs. Stim 5' | ** | 0.0080 | |
| Rest vs. Recov 5' (Stim 1') | *** | < 0.0001 | |
| Stim 1' vs. Stim 5' | ns | > 0.9999 | |
| Stim 1' vs. Recov 5' (Stim 1') | *** | < 0.0001 | |
| Stim 5' vs. Recov 5' (Stim 1') | *** | < 0.0001 | |

**Figure 3B.**

| Genotype/Condition | Mean ± s.e.m. | N | n |
|---|---|---|---|
| Rest | 100 ± 7 % | 7 | 122 |
| Stim 1' | 442 ± 28 % | 8 | 141 |
| Recov 5' (Stim 1') | 178 ± 7 % | 6 | 112 |

| Compared group | Statistical significance | *P*-value | Statistical test |
|---|---|---|---|
| ***PLA Otoferlin/PKCα:*** | | | |
| Rest vs. Stim 1' | *** | < 0.0001 | Kruskal-Wallis test followed by Dunn's multiple comparison test |
| Rest vs. Recov 5' (Stim 1') | *** | < 0.0001 | |
| Stim 1' vs. Recov 5' (Stim 1') | *** | < 0.0001 | |

**Figures 4B-C.**

| Genotype/Condition | Mean ± s.e.m. | | N | n |
|---|---|---|---|---|
| | Immunofluorescence | Apical/basal ratio | | |
| WT | 100 ± 2 % | 1.06 ± 0.03 | 6 | 233 |
| *Otof* $^{-/-}$ | 92 ± 1 % | 0.66 ± 0.02 | 6 | 205 |

| Compared group | Statistical significance | *P*-value | Statistical test |
|---|---|---|---|
| ***PKC$\alpha$ immunofluorescence:*** | | | |
| WT vs. *Otof* $^{-/-}$ | ** | 0.0054 | Mann-Whitney two-tailed t-test |
| ***Apical/basal PKC$\alpha$ ratio:*** | | | |
| WT vs. *Otof* $^{-/-}$ | *** | < 0.0001 | Mann-Whitney two-tailed t-test |

**Figure 4G.**

| Genotype/Condition | Mean ± s.e.m. | N | n |
|---|---|---|---|
| WT Rest | 1.05 ± 0.03 | 6 | 150 |
| WT Stim 1' | 0.76 ± 0.02 | 7 | 179 |
| WT Stim 5' | 0.85 ± 0.06 | 1 | 37 |
| WT Recov 5' (Stim 1') | 1.32 ± 0.04 | 6 | 195 |
| *Otof* $^{-/-}$ Rest | 1.15 ± 0.12 | 1 | 15 |
| *Otof* $^{-/-}$ Stim 1' | 0.75 ± 0.03 | 2 | 50 |
| *Otof* $^{-/-}$ Stim 5' | 0.81 ± 0.03 | 2 | 46 |
| *Otof* $^{-/-}$ Recov 5' (Stim 1') | 0.83 ± 0.04 | 1 | 26 |

| Compared group | Statistical significance | *P*-value | Statistical test |
|---|---|---|---|
| ***WT vs. Otof $^{-/-}$ apical/basal PKC$\alpha$ ratio:*** | | | |
| WT Rest vs. *Otof* $^{-/-}$ Rest | ns | > 0.9999 | Kruskal-Wallis test followed by Dunn's multiple comparison test |
| WT Rest vs. WT Stim 1' | *** | < 0.0001 | |
| WT Rest vs. *Otof* $^{-/-}$ Stim 1' | *** | < 0.0001 | |
| WT Rest vs. WT Stim 5' | * | 0.0101 | |
| WT Rest vs. *Otof* $^{-/-}$ Stim 5' | ** | 0.0051 | |
| WT Rest vs. WT Recov 5' (Stim 1') | *** | < 0.0001 | |
| WT Rest vs. *Otof* $^{-/-}$ Recov 5' (Stim 1') | ns | 0.1708 | |
| *Otof* $^{-/-}$ Rest vs. WT Stim 1' | ** | 0.0011 | |
| *Otof* $^{-/-}$ Rest vs. *Otof* $^{-/-}$ Stim 1' | ** | 0.0023 | |
| *Otof* $^{-/-}$ Rest vs. WT Stim 5' | ns | 0.0754 | |
| *Otof* $^{-/-}$ Rest vs. *Otof* $^{-/-}$ Stim 5' | ns | 0.0740 | |
| *Otof* $^{-/-}$ Rest vs. WT Recov 5' (Stim 1') | ns | > 0.9999 | |
| *Otof* $^{-/-}$ Rest vs. *Otof* $^{-/-}$ Recov 5' (Stim 1') | ns | 0.2537 | |
| WT Stim 1' vs. *Otof* $^{-/-}$ Stim 1' | ns | > 0.9999 | |
| WT Stim 1' vs. WT Stim 5' | ns | > 0.9999 | |
| WT Stim 1' vs. *Otof* $^{-/-}$ Stim 5' | ns | > 0.9999 | |
| WT Stim 1' vs. WT Recov 5' (Stim 1') | *** | < 0.0001 | |
| WT Stim 1' vs. *Otof* $^{-/-}$ Recov 5' (Stim 1') | ns | > 0.9999 | |

| | | |
|---|---|---|
| *Otof*$^{-/-}$ Stim 1' vs. WT Stim 5' | ns | > 0.9999 |
| *Otof*$^{-/-}$ Stim 1' vs. *Otof*$^{-/-}$ Stim 5' | ns | > 0.9999 |
| *Otof*$^{-/-}$ Stim 1' vs. WT Recov 5' (Stim 1') | *** | < 0.0001 |
| *Otof*$^{-/-}$ Stim 1' vs. *Otof*$^{-/-}$ Recov 5' (Stim 1') | ns | > 0.9999 |
| WT Stim 5' vs. *Otof*$^{-/-}$ Stim 5' | ns | > 0.9999 |
| WT Stim 5' vs. WT Recov 5' (Stim 1') | *** | < 0.0001 |
| WT Stim 5' vs. *Otof*$^{-/-}$ Recov 5' (Stim 1') | ns | > 0.9999 |
| *Otof*$^{-/-}$ Stim 5' vs. WT Recov 5' (Stim 1') | *** | < 0.0001 |
| *Otof*$^{-/-}$ Stim 5' vs. *Otof*$^{-/-}$ Recov 5' (Stim 1') | ns | > 0.9999 |
| WT Recov 5' (Stim 1') vs. *Otof*$^{-/-}$ Recov 5' (Stim 1') | *** | < 0.0001 |

**Figure 5B.**

| Genotype/Condition | Mean ± s.e.m. | N | n |
|---|---|---|---|
| Rest | 100 ± 11 % | 3 | 100 |
| Stim 1’ | 234 ± 13 % | 1 | 37 |
| Stim 5' | 438 ± 38 % | 2 | 52 |
| BIM I + Stim 5' | 122 ± 2 % | 1 | 34 |
| BIM I+KN-93 + Stim 5' | 97 ± 4 % | 1 | 50 |
| PMA 1’ | 77 ± 7 % | 2 | 61 |
| PMA 5’ | 157 ± 4 % | 2 | 66 |
| PMA 15’ | 139 ± 6 % | 2 | 48 |

| Compared group | Statistical significance | *P*-value | Statistical test |
|---|---|---|---|
| ***PLA Otoferlin/P-Serine:*** | | | |
| Rest vs. Stim 1' | *** | < 0.0001 | Kruskal-Wallis test followed by Dunn’s multiple comparison test |
| Rest vs. Stim 5' | *** | < 0.0001 | |
| Stim 1' vs. Stim 5' | ns | > 0.9999 | |
| Stim 5' vs. BIM I + Stim 5' | ** | 0.0013 | |
| Stim 5' vs. BIM I+KN-93 + Stim 5' | *** | < 0.0001 | |
| BIM I + Stim 5' vs. BIM I+KN-93 + Stim 5' | * | 0.0250 | |
| Rest vs. BIM I + Stim 5' | * | 0.0133 | |
| Rest vs. BIM I+KN-93 + Stim 5' | ns | > 0.9999 | |
| Rest vs. PMA 1' | ns | 0.3464 | |
| Rest vs. PMA 5' | *** | < 0.0001 | |
| Rest vs. PMA 15' | *** | 0.0002 | |
| PMA 1' vs. PMA 5' | *** | < 0.0001 | |
| PMA 1' vs. PMA 15' | *** | < 0.0001 | |
| PMA 5' vs. PMA 15' | ns | 0.1802 | |

**Figure 6B.**

| Genotype/Condition | Mean ± s.e.m. | N | n |
|---|---|---|---|
| Rest | 100 ± 5 % | 2 | 43 |
| Stim 1' | 138 ± 3 % | 2 | 44 |
| Recov 5' (Stim 1') | 107 ± 3 % | 1 | 23 |

| Compared group | Statistical significance | *P*-value | Statistical test |
|---|---|---|---|
| ***Myosin VI immunofluorescence:*** | | | |
| Rest vs. Stim 1' | *** | < 0.0001 | Kruskal-Wallis test followed by Dunn's multiple comparison test |
| Rest vs. Recov 5' (Stim 1') | ns | > 0.9999 | |
| Stim 1' vs. Recov 5' (Stim 1') | *** | < 0.0001 | |

**Figure 6D.**

| Genotype/Condition | Mean ± s.e.m. | N | n |
|---|---|---|---|
| Rest | 100 ± 3 % | 6 | 265 |
| Stim 1' | 173 ± 4 % | 3 | 170 |
| Recov 5' (Stim 1') | 183 ± 4 % | 3 | 153 |
| BIM I + Stim 1' | 102 ± 6 % | 1 | 37 |
| PMA 5' | 133 ± 3 % | 3 | 122 |
| PMA 15' | 149 ± 9 % | 3 | 96 |

| Compared group | Statistical significance | *P*-value | Statistical test |
|---|---|---|---|
| ***PLA Otoferlin/Myosin VI:*** | | | |
| Rest vs. Stim 1' | *** | < 0.0001 | Kruskal-Wallis test followed by Dunn's multiple comparison test |
| Rest vs. Recov 5' (Stim 1') | *** | < 0.0001 | |
| Rest vs. BIM I + Stim 1' | ns | > 0.9999 | |
| Rest vs. PMA 5' | *** | < 0.0001 | |
| Rest vs. PMA 15' | *** | < 0.0001 | |
| Stim 1' vs. Recov 5' (Stim 1') | ns | > 0.9999 | |
| Stim 1' vs. BIM I + Stim 1' | *** | < 0.0001 | |
| Stim 1' vs. PMA 5' | *** | < 0.0001 | |
| Stim 1' vs. PMA 15' | *** | < 0.0001 | |
| Recov 5' (Stim 1') vs. BIM I + Stim 1' | *** | < 0.0001 | |
| Recov 5' (Stim 1') vs. PMA 5' | *** | < 0.0001 | |
| Recov 5' (Stim 1') vs. PMA 15' | *** | < 0.0001 | |
| BIM I + Stim 1' vs. PMA 5' | ** | 0.0013 | |
| BIM I + Stim 1' vs. PMA 15' | *** | 0.0002 | |
| PMA 5' vs. PMA 15' | ns | > 0.9999 | |

**Figure 6F-G.**

| Genotype/Condition | Mean ± s.e.m. | | N | n |
|---|---|---|---|---|
| | Immunofluorescence | Apical/basal ratio | | |
| Rest | 100 ± 2 % | 1.12 ± 0.06 | 3 | 89 |
| Stim 1' | 163 ± 8 % | 0.83 ± 0.04 | 3 | 92 |
| Recov 5' (Stim 1') | 118 ± 2 % | 1.19 ± 0.08 | 3 | 71 |

| Compared group | Statistical significance | *P*-value | Statistical test |
|---|---|---|---|
| ***Vglut3 immunofluorescence:*** | | | |
| Rest vs. Stim 1' | *** | < 0.0001 | Kruskal-Wallis test followed by Dunn's multiple comparison test |
| Rest vs. Recov 5' (Stim 1') | *** | < 0.0001 | |
| Stim 1' vs. Recov 5' (Stim 1') | ns | > 0.9999 | |
| ***Apical/basal Vglut3 ratio:*** | | | |
| Rest vs. Stim 1' | *** | 0.0005 | Kruskal-Wallis test followed by Dunn's multiple comparison test |
| Rest vs. Recov 5' (Stim 1') | ns | > 0.9999 | |
| Stim 1' vs. Recov 5' (Stim 1') | *** | 0.0009 | |

**Figure 6I.**

| Genotype/Condition | Mean ± s.e.m. | N | n |
|---|---|---|---|
| Rest | 100 ± 2 % | 5 | 78 |
| Stim 1' | 104 ± 1 % | 4 | 146 |
| Recov 5' (Stim 1') | 95 ± 2 % | 3 | 93 |

| Compared group | Statistical significance | *P*-value | Statistical test |
|---|---|---|---|
| ***PLA Otoferlin/Vglut3:*** | | | |
| Rest vs. Stim 1' | ns | 0.0961 | Kruskal-Wallis test followed by Dunn's multiple comparison test |
| Rest vs. Recov 5' (Stim 1') | ns | 0.5761 | |
| Stim 1' vs. Recov 5' (Stim 1') | *** | 0.0005 | |

**Figure 7B-C.**

| Genotype/Condition | Mean ± s.e.m. | | N | n |
|---|---|---|---|---|
| | Immunofluorescence | Apical/basal ratio | | |
| Rest | 100 ± 1 % | 1.04 ± 0.02 | 15 | 296 |
| Stim 1' | 62 ± 2 % | 0.84 ± 0.03 | 11 | 174 |
| Recov 5' (Stim 1') | 73 ± 2 % | 1.07 ± 0.05 | 8 | 141 |
| BIM I + Stim 1' | 99 ± 3 % | 1.14 ± 0.07 | 1 | 26 |

| Compared group | Statistical significance | *P*-value | Statistical test |
|---|---|---|---|
| ***Calbindin immunofluorescence:*** | | | |
| Rest vs. Stim 1' | *** | < 0.0001 | Kruskal-Wallis test followed by Dunn's multiple comparison test |
| Rest vs. Recov 5' (Stim 1') | *** | < 0.0001 | |
| Rest vs. BIM I + Stim 1' | ns | > 0.9999 | |
| Stim 1' vs. Recov 5' (Stim 1') | * | 0.0152 | |
| Stim 1' vs. BIM I + Stim 1' | *** | < 0.0001 | |
| Recov 5' (Stim 1') vs. BIM I + Stim 1' | *** | < 0.0001 | |
| ***Apical/basal Calbindin ratio:*** | | | |
| Rest vs. Stim 1' | *** | < 0.0001 | Kruskal-Wallis test followed by Dunn's multiple comparison test |
| Rest vs. Recov 5' (Stim 1') | ns | 0.6380 | |
| Rest vs. BIM I + Stim 1' | ns | 0.8940 | |
| Stim 1' vs. Recov 5' (Stim 1') | *** | 0.0007 | |
| Stim 1' vs. BIM I + Stim 1' | *** | 0.0001 | |
| Recov 5' (Stim 1') vs. BIM I + Stim 1' | ns | 0.1856 | |

**Figure 7D**.

| Genotype/Condition | Mean ± s.e.m. | | N | n |
|---|---|---|---|---|
| | Calbindin | Otoferlin | | |
| WT | 100 ± 1 % | 100 ± 1 % | 8 | 176 |
| $Otof^{I515T/I515T}$ | 72 ± 3 % | 42 ± 1 % | 3 | 83 |
| $Otof^{Pga/Pga}$ | 92 ± 2 % | 30 ± 1 % | 4 | 76 |
| $Otof^{+/-}$ | 83 ± 3 % | 51 ± 2 % | 3 | 99 |
| $Otof^{-/-}$ | 56 ± 1 % | 0 ± 0 % | 4 | 108 |

| Compared group | Statistical significance | *P*-value | Statistical test |
|---|---|---|---|
| ***Calbindin levels:*** | | | |
| WT vs. $Otof^{I515T/I515T}$ | *** | < 0.0001 | Kruskal-Wallis test followed by Dunn's multiple comparison test |
| WT vs. $Otof^{Pga/Pga}$ | ns | 0.5900 | |
| WT vs. $Otof^{+/-}$ | *** | 0.0002 | |
| WT vs. $Otof^{-/-}$ | *** | < 0.0001 | |
| $Otof^{I515T/I515T}$ vs. $Otof^{Pga/Pga}$ | * | 0.0124 | |
| $Otof^{I515T/I515T}$ vs. $Otof^{+/-}$ | ns | > 0.9999 | |
| $Otof^{I515T/I515T}$ vs. $Otof^{-/-}$ | * | 0.0186 | |
| $Otof^{Pga/Pga}$ vs. $Otof^{+/-}$ | ns | > 0.9999 | |
| $Otof^{Pga/Pga}$ vs. $Otof^{-/-}$ | *** | < 0.0001 | |
| $Otof^{+/-}$ vs. $Otof^{-/-}$ | *** | < 0.0001 | |
| ***Otoferlin levels:*** | | | |
| WT vs. $Otof^{I515T/I515T}$ | *** | < 0.0001 | Kruskal-Wallis test followed by Dunn's multiple comparison test |
| WT vs. $Otof^{Pga/Pga}$ | *** | < 0.0001 | |
| WT vs. $Otof^{+/-}$ | *** | < 0.0001 | |
| WT vs. $Otof^{-/-}$ | *** | < 0.0001 | |
| $Otof^{I515T/I515T}$ vs. $Otof^{Pga/Pga}$ | ns | 0.4603 | |
| $Otof^{I515T/I515T}$ vs. $Otof^{+/-}$ | ns | > 0.9999 | |

| | | |
|---|---|---|
| *Otof*$^{I515T/I515T}$ vs. *Otof*$^{-/-}$ | *** | < 0.0001 |
| *Otof*$^{Pga/Pga}$ vs. *Otof*$^{+/-}$ | *** | 0.0006 |
| *Otof*$^{Pga/Pga}$ vs. *Otof*$^{-/-}$ | *** | < 0.0001 |
| *Otof*$^{+/-}$ vs. *Otof*$^{-/-}$ | *** | < 0.0001 |

**Figure 7E**.

| Genotype/Condition | Mean ± s.e.m. | | N | n |
|---|---|---|---|---|
| | Calbindin | Otoferlin | | |
| WTB6 - AAV | 100 ± 2 % | 100 ± 1 % | 11 | 276 |
| WTCD1B6F1 + AAV.eGFP | 94 ± 3 % | 104 ± 4 % | 8 | 168 |
| WTCD1B6F1 + DualAAV-TS | 117 ± 4 % | 147 ± 5 % | 3 | 62 |
| *Otof*$^{-/-}$ CD1B6F1 + DualAAV-TS | 70 ± 5 % | 31 ± 3 % | 1 | 13 |
| *Otof*$^{-/-}$ CD1B6F1 + DualAAV-Hyb | 63 ± 3 % | 29 ± 2 % | 5 | 64 |
| *Otof*$^{-/-}$ CD1B6F1 - AAV | 50 ± 2 % | 3 ± 0 % | 6 | 142 |

| Compared group | Statistical significance | *P*-value | Statistical test |
|---|---|---|---|
| ***Calbindin levels:*** | | | |
| WTB6 – AAV vs. WTCD1B6F1 + AAV.eGFP | ns | 0.36 | Kruskal-Wallis test followed by Dunn's multiple comparison test |
| WTB6 – AAV vs. WTCD1B6F1 + DualAAV-TS | ** | 0.002 | |
| WTB6 – AAV vs. *Otof*$^{-/-}$ CD1B6F1 + DualAAV-TS | *** | < 0.0001 | |
| WTB6 – AAV vs. *Otof*$^{-/-}$ CD1B6F1 + DualAAV-Hyb | *** | < 0.0001 | |
| WTB6 – AAV vs. *Otof*$^{-/-}$CD1B6F1 - AAV | *** | < 0.0001 | |
| *Otof*$^{-/-}$ CD1B6F1 + DualAAV-TS vs. *Otof*$^{-/-}$ CD1B6F1 + DualAAV-Hyb | ns | 0.38 | |
| *Otof*$^{-/-}$ CD1B6F1 + DualAAV-TS vs. *Otof*$^{-/-}$ CD1B6F1 - AAV | ** | 0.0095 | |
| *Otof*$^{-/-}$ CD1B6F1 + DualAAV-Hyb vs. *Otof*$^{-/-}$ CD1B6F1 – AAV | * | 0.0292 | |
| ***Otoferlin levels:*** | | | |
| WTB6 – AAV vs. WTCD1B6F1 + AAV.eGFP | ns | 0.58 | Kruskal-Wallis test followed by Dunn's multiple comparison test |
| WTB6 – AAV vs. WTCD1B6F1 + DualAAV-TS | *** | < 0.0001 | |
| WTB6 – AAV vs. *Otof*$^{-/-}$ CD1B6F1 + DualAAV-TS | *** | < 0.0001 | |
| WTB6 – AAV vs. *Otof*$^{-/-}$ CD1B6F1 + DualAAV-Hyb | *** | < 0.0001 | |
| WTB6 – AAV vs. *Otof*$^{-/-}$ CD1B6F1 - AAV | *** | < 0.0001 | |
| *Otof*$^{-/-}$ CD1B6F1 + DualAAV-TS vs. *Otof*$^{-/-}$ CD1B6F1 + DualAAV-Hyb | ns | > 0.9999 | |

| | | |
|---|---|---|
| *Otof*$^{-/-}$CD1B6F1 + DualAAV-TS vs. *Otof*$^{-/-}$ CD1B6F1 - AAV | *** | < 0.0001 |
| *Otof*$^{-/-}$CD1B6F1 + DualAAV-Hyb vs. *Otof*$^{-/-}$CD1B6F1 - AAV | *** | < 0.0001 |

**Figure 7H.**

| Genotype/Condition | Mean ± s.e.m. | N | n |
|---|---|---|---|
| Rest | 100 ± 2 % | 7 | 327 |
| Stim 1' | 560 ± 26 % | 4 | 168 |
| Recov 5' (Stim 1') | 77 ± 5 % | 2 | 107 |
| BIM I + Stim 1' | 101 ± 3 % | 2 | 98 |
| PMA 5’ | 175 ± 4 % | 3 | 114 |
| PMA 15’ | 161 ± 3 % | 3 | 127 |

| Compared group | Statistical significance | *P*-value | Statistical test |
|---|---|---|---|
| ***PLA Otoferlin/Calbindin:*** | | | |
| Rest vs. Stim 1' | *** | < 0.0001 | Kruskal-Wallis test followed by Dunn’s multiple comparison test |
| Rest vs. Recov 5' (Stim 1') | *** | < 0.0001 | |
| Rest vs. PMA 5' | *** | < 0.0001 | |
| Rest vs. PMA 15' | *** | < 0.0001 | |
| Rest vs. BIM I + Stim 1' | ns | > 0.9999 | |
| Stim 1' vs. Recov 5' (Stim 1') | *** | < 0.0001 | |
| Stim 1' vs. PMA 5' | *** | < 0.0001 | |
| Stim 1' vs. PMA 15' | *** | < 0.0001 | |
| Stim 1' vs. BIM I + Stim 1' | *** | < 0.0001 | |
| Recov 5' (Stim 1') vs. PMA 5' | *** | < 0.0001 | |
| Recov 5' (Stim 1') vs. PMA 15' | *** | < 0.0001 | |
| Recov 5' (Stim 1') vs. BIM I + Stim 1' | *** | 0.0006 | |
| PMA 5' vs. PMA 15' | ns | > 0.9999 | |
| PMA 5' vs. BIM I + Stim 1' | *** | < 0.0001 | |
| PMA 15' vs. BIM I + Stim 1' | *** | < 0.0001 | |

**Figure EV4.**

| Genotype/Condition | Mean ± s.e.m. | N | n |
|---|---|---|---|
| Rest | 100 ± 6 % | 2 | 75 |
| Stim 1' | 158 ± 5 % | 2 | 94 |
| Recov 5' (Stim 1') | 82 ± 6 % | 2 | 61 |

| Compared group | Statistical significance | *P*-value | Statistical test |
|---|---|---|---|
| ***PLA PKC$\alpha$/Calbindin:*** | | | |
| Rest vs. Stim 1' | *** | < 0.0001 | Kruskal-Wallis test followed by Dunn’s multiple comparison test |
| Rest vs. Recov 5' (Stim 1') | ** | 0.0051 | |
| Stim 1' vs. Recov 5' (Stim 1') | *** | < 0.0001 | |

#### Appendix Table S2. Prediction of PKC phosphorylation sites in otoferlin.

| Tool | Model | Sites | | Reference |
|---|---|---|---|---|
| **NetPhos 3.1** | ANN | T24; S55; S74; S173; S224; T229; S230; T285; S319; T342; T466; S467; S530; S541; S563; S683; S715; S784; T904; T954; T1050; S1224; T1457; T1504; T1538; T1577; T1597; S1646; T1840; S1859; T1940 | http://www.cbs.dtu.dk/services/NetPhos-3.1/ | (Blom *et al*, 2004) |
| **KinasePhos 1.0** | HMM | T9; S74; S158; T159; S162; S166; S183; S185; S237; S246; T285; T331; T446; S467; S520; S541; S563; S685; T750; S775; S784; S785; T787; S810; T819; S866; T904; S970; T1083; S1130; S1223; S1224; T1242; T1261; S1293; T1318; T1416; T1424; T1482; T1538; S1566; T1577; T1597; T1624; T1639; T1756; S1789; S1796; T1809 | http://kinasephos.mbc.nctu.edu.tw/ | (Huang *et al*, 2005) |
| **PSSP** | BDT | T24; S29; S55; S74; S113; S145; S158; S162; S166; S185; S233; S237; S246; S286; S290; S335; S434; S467; S472; S501; S520; S541; S563; S753; S775; S784; S785; S796; S803; S810; T819; T826; S918; S1099; S1130; S1223; S1224; S1293; S1351; S1355; S1436; S1566; S1579; S1705; S1789; S1793; T1840; S1933 | http://ppsp.biocuckoo.org/ | (Xue *et al*, 2006) |
| **GPS 3.0** | PSSM, GA | T24; S55; S74; S158; S162; S166; S185; S219; S233; S467; S775; S777; S784; S785; S796; S810; T819; S918; S1040; S1099; S1224; S1351; S1355; S1965 | http://gps.biocuckoo.org/online.php | (Xue *et al*, 2011) |

Data information: Mouse otoferlin isoform 4 (NCBI accession number NP_001300696.1) was used for predictions. ANN, artificial neural network; HMM, Hidden Markov Models; PSSM, position-specific scoring matrices; GA, genetic algorithm; BDT, Bayesian decision theory.
